# Chronic Ca^2+^ imaging of cortical neurons with long-term expression of GCaMP-X

**DOI:** 10.1101/2022.01.09.475579

**Authors:** Jinli Geng, Wenxiang Li, Yingjun Tang, Yunming Gao, Yitong Lu, Ping Li, Xiaomei Li, Ping Wang, Yubo Fan, Yaxiong Yang, Zengcai V. Guo, Xiaodong Liu

## Abstract

Dynamic Ca^2+^ signals reflect acute changes in membrane excitability (e.g. sensory response), and also mediate intracellular signaling cascades normally of longer time scales (e.g., Ca^2+^-dependent neuritogenesis). In both cases, chronic Ca^2+^ imaging has been often desired, but largely hindered by unexpected cytotoxicity intrinsic to GCaMP, a popular series of genetically-encoded Ca^2+^ indicators. Here, we demonstrate that the recently developed GCaMP-X outperforms GCaMP in long-term probe expression and/or chronic Ca^2+^ imaging. GCaMP-X shows much improved compatibility with neurons and thus more reliable than GCaMP as demonstrated *in vivo* by acute Ca^2+^ responses to whisker deflection or spontaneous Ca^2+^ fluctuations over an extended time frame. Chronic Ca^2+^ imaging data (*≥*1 month) are acquired from the same set of cultured cortical neurons, unveiling that spontaneous/local Ca^2+^ activities would progressively develop into autonomous/global Ca^2+^ oscillations. Besides the morphological indices of neurite length or soma size, the major metrics of oscillatory Ca^2+^, including rate, amplitude, synchrony among different neurons or organelles have also been examined along with the developmental stages. Both neuritogenesis and Ca^2+^ signals are dysregulated by GCaMP in virus-infected or transgenic neurons, in direct contrast to GCaMP-X without any noticeable side-effect. Such *in vitro* data altogether consolidate the unique importance of oscillatory Ca^2+^ to activity-dependent neuritogenesis, as one major factor responsible for the distinctions between GCaMP vs GCaMP-X *in vivo*. For the first time with GCaMP-X of long-term expression in neurons, spontaneous and sensory-evoked Ca^2+^ activities are imaged and evaluated both *in vitro* and *in vivo*, providing new opportunities to monitor neural development or other chronic processes concurrently with Ca^2+^ dynamics.

## Introduction

Ca^2+^ signals play pivotal roles in the brain, closely involved in membrane excitability, sensory transduction, synaptic transmission, neural development and plasticity (Berridge et al., 2003). Ca^2+^ dysregulations are linked with the mental disorders including Parkinson’s diseases, Alzheimer’s diseases, epilepsy and schizophrenia (Chan et al., 2007; Fernandez de Sevilla et al., 2006; Khan et al., 2020; Liebscher et al., 2016), suggesting that multiple factors and cascades may converge to Ca^2+^ as one of the central factors underlying brain diseases, referred as the ‘calcium hypothesis’ (Berridge, 2010). According to their downstream consequences, cellular Ca^2+^ signals could be categorized as genomic versus non-genomic, to reflect the fact that in some cases where gene expressions are regulated (chronic) versus in other cases only the (acute) functions of existing proteins are concerned.

Acutely, cellular Ca^2+^ is able to indicate the activities of single neurons, such as spontaneous fluctuations and stimulus-evoked responses (Chen et al., 2013; O’Banion and Yasuda, 2020). Ca^2+^ imaging is often utilized to measure neuronal excitability, as one alternative to electrical recording. In fact, genetically encoded Ca^2+^ indicators (GECI) represented by GCaMP (based on calmodulin or CaM, and Ca^2+^/CaM-binding motif M13) have been broadly applied to monitor neurons and other excitable cells (Akerboom et al., 2012; Chen et al., 2013; Dana et al., 2019; Nakai et al., 2001; Tallini et al., 2006; Tian et al., 2009; Yang et al., 2018). In addition to a faithful indicator of acute responses (in the timescale of seconds/minutes, such as a burst of action potentials), Ca^2+^ is often tightly coupled to various chronic effects or processes, e.g., Ca^2+^-dependent gene transcription and expression, neurite outgrowth or pruning, long-term potentiation or depression, learning and memory, and neural degeneration (O’Banion and Yasuda, 2020). Therefore, it is highly desirable to monitor the long-term Ca^2+^ dynamics (days/weeks or longer) for cells, tissues, organs or even whole organisms, which would greatly facilitate mechanistic understanding of diverse pathophysiology (Garcia et al., 2017). In fact, even for acute Ca^2+^ responses, long-term imaging lasting multiple hours or days is also often in need, e.g., to examine the plastic changes of responses and the underlying cellular/circuitry mechanisms (Schuette et al., 2020).

However, when the expression level of GCaMP is being increased with time in neurons, the cytotoxicity becomes more and more apparent as demonstrated in previous reports including ours (Yang et al., 2018). Consequently, all the aforementioned applications (especially chronic imaging) would be adversely affected, since GCaMP side-effects are accumulated and exacerbated with time. Expression of GECI is not acute by default because it takes hours or even days for GECI to get expressed in cells, which would make the side-effects inevitable. However, such effects are not always apparent especially when the absolute level and/or the time course of GCaMP expression are intentionally limited. For instance, when employing viral infection, imaging experiments are restricted within the empirical time-window to avoid nuclear GCaMP accumulation; or for transgenic mice, special promoters, conditional expression and other tactics are utilized to prevent GCaMP overexpression (Madisen et al., 2015). With such work-around solutions, a great amount of new knowledge has been achieved, particularly from *in vivo* imaging of live animals. Until recently, the side-effects are unveiled to arise from GCaMP perturbations on apoCaM signaling, represented by L-type Ca^2+^ channels dysregulated by GCaMP of long-term and/or high-level expression (Steinmetz et al., 2017; Tian et al., 2009; Yang et al., 2018). Therefore, the side-effects of GCaMP have seriously jeopardized the quality and impacts that could have been achieved by numerous GCaMP applications. As demonstrated by this work, GCaMP cyto-toxicities are indeed present and persistent in chronic imaging experiments under all the conditions we have tested: either with old or new versions of GCaMP, either for viral or transgenic expression of GCaMP, either *in vitro* (cultured neurons) or *in vivo* (living mice), and either monitoring acute (sensory responses) or chronic (spontaneous oscillations) activities.

In this work, GCaMP-X with enhanced neuron compatibility (Yang et al., 2018) was applied to different scenarios including cultured, virus-infected or transgenic neurons in *vitro* and *in vivo*. GCaMP-X imaging *in vivo* is able to faithfully capture the acute sensory responses during the extended period of time beyond the “safe window” (>8 weeks) of GCaMP. In contrast, Ca^2+^ signals are impaired for neurons expressing GCaMP including those from transgenic mice, suggesting scrutiny is required in neuron selection and data interpretation. Importantly, by focusing on spontaneous Ca^2+^ activities including autonomous Ca^2+^ oscillations, chronic GCaMP-X imaging has been successfully implemented *in vitro* and *in vivo*, featured with long-term overexpression (free of CaM-interference), high spatiotemporal contents (multiple weeks and intact neuronal network) and subcellular resolution (cytosolic versus nuclear), all of which are nearly infeasible if using conventional GCaMP.

## RESULTS

### Design principles of GCaMP-X validated with newer versions of GCaMP

Recently, GCaMP has been updated to its newest versions jGCaMP7 (but also see the bioRxiv preprint of jGCaMP8 (Zhang et al., 2021)), with enhanced sensing performance of multiple aspects over the previous GCaMP6 (Dana et al., 2019; Grødem et al., 2021). Considering that the design of jGCaMP7 is also on the basis of CaM, we postulated that GCaMP-contained apoCaM would still be problematic. We chose jGCaMP7b to further validate the design principles of neuron-compatible GCaMP-X previously established with GCaMP6 and earlier versions (Yang et al., 2018). Following the protocol of transient transfection (***Figure 1A***), jGCaMP7b accumulated into the nuclei indexed by N/C (nuclear/cytosolic) ratio in a substantial subpopulation of cortical neurons (***Figure 1B***), one major feature of GCaMP side-effects. Notably, jGCaMP7b exhibited even more severe nuclear accumulation than other GCaMP variants, which may account for the nuclear jGCaMP7b evidenced *in vivo* (Dana et al., 2019). Accordingly, the total length (***Figure 1C***) and the complexity (***Figure 1D***) of neurites were significantly reduced in jGCaMP7b-expressing neurons. The apoCaM-binding motif (CBM) and the localization tags were then appended onto jGCaMP7b, following the design of GCaMP-X (Yang et al., 2018), to construct jGCaMP7b-X_C_ (cytosolic Ca^2+^) and jGCaMP7b-X_N_ (nuclear Ca^2+^). GCaMP-X is supposed to eliminate the cytotoxicity intrinsic to GCaMP, thus capable of long-term Ca^2+^ monitoring. Depicted by neurite tracing (***Figure 1A***), both cytosolic and nuclear versions of GCaMP7-X have greatly enhanced the compatibility with neurons. In fact, neurons expressing GCaMP7-X are essentially indistinguishable from GFP control neurons, in direct contrast to GCaMP7 (***Figure 1C, D***). To further confirm the above advantages of GCaMP7-X over GCaMP7 in an extended timeframe (up to DIV 42), adeno-associated viruses (AAV) were applied to cortical neurons for side-by-side comparison (***Figure 1—figure supplement 1***). Similar conclusions have been reached here that GCaMP7-X well conserves the general health of neurons whereas GCaMP7 causes progressive damages to neurons. In light of GCaMP perturbations on apoCaM signaling (Yang et al., 2018), we examined the effects of GCaMP7b on recombinant CaV1 channels in HEK293 cells. jGCaMP7b significantly altered the major properties of CaV1.3 gating, i.e., abnormally stronger calcium-dependent inactivation (CDI) and larger voltage-gated activation (VGA) (***Figure 1E***). Collectively, the mechanisms and design principles established with GCaMP6 and earlier versions are readily extendable onto CaM-based GECIs, as validated with jGCaMP7.

**Figure 1.**
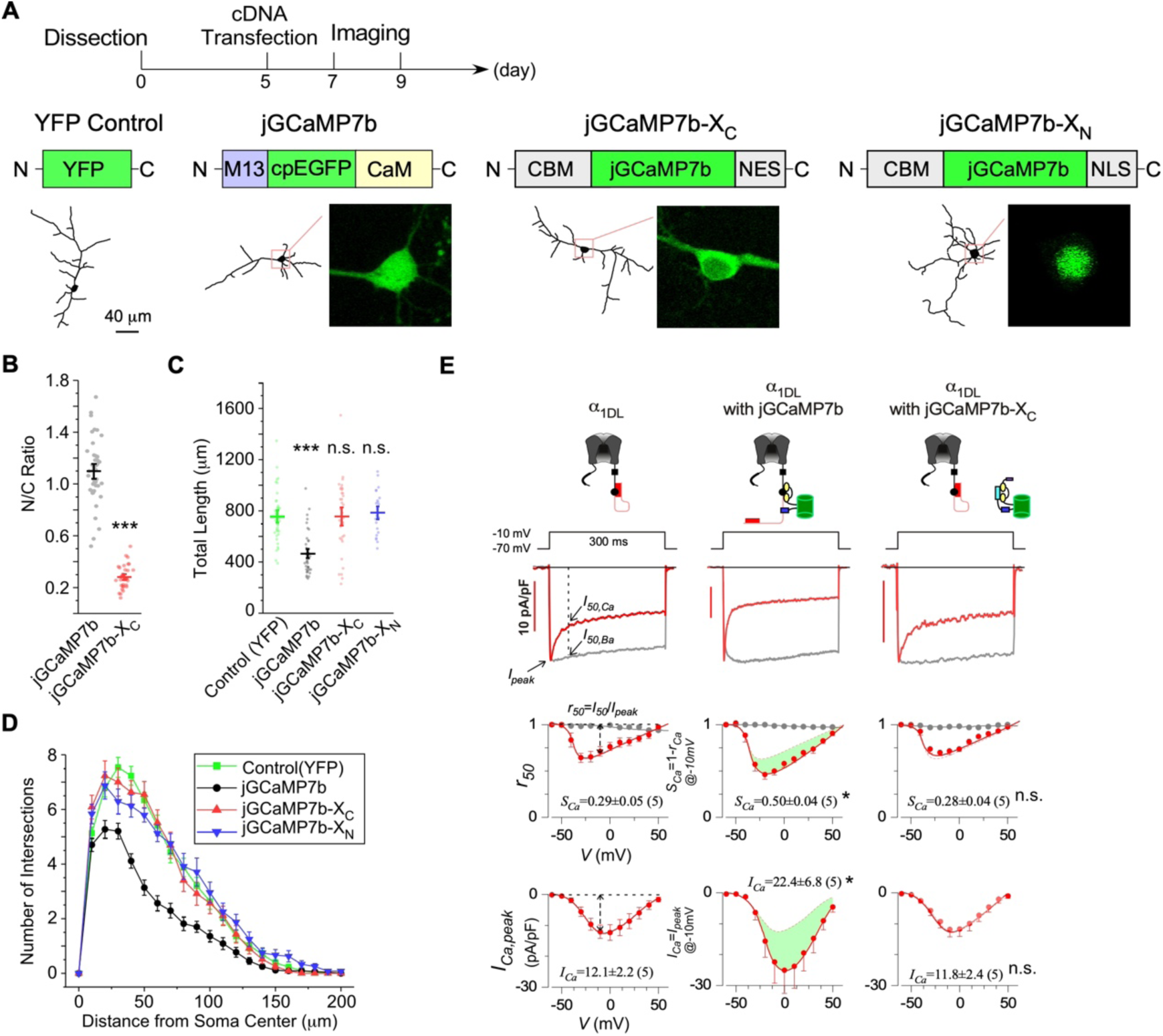
The design principles extendable to jGCaMP7 and jGCaMP7-X. (**A**) Cultured cortical neurons from newborn mice were transiently transfected with YFP, jGCaMP7b, jGCaMP7b-X_C_ or jGCaMP7b-X_N_, respectively on DIV 5-7, then imaged by confocal microscopy on DIV 7-9. As illustrated, apo<u>C</u>aM <u>b</u>inding <u>m</u>otif (CBM) was fused onto N-terminus of GCaMP and the tags of localization signals (<u>n</u>uclear <u>e</u>xport <u>s</u>ignal or <u>n</u>uclear localization <u>s</u>ignal, NES/NLS) were fused to the C-terminus of GCaMP to construct GCaMP-X_C_ or GCaMP-X_N_, respectively. Neurite tracing and subcellular GCaMP distributions are shown below. (**B**) N/C ratio of jGCaMP7b or jGCaMP7b-X_C_ in neurons, by calculating the nucleus versus cytosol ratio of fluorescence intensities. (**C, D**) Total length (**C**) and *Sholl* analysis (**D**) for cortical neurons expressing YFP or GCaMP variants. (**E**) Electrophysiological validations of jGCaMP7b versus jGCaMP7b-X with recombinant CaV1.3 channels. Full-length CaV1.3 channels (*α*_1DL_) were expressed in HEK293 cells alone (left) or with jCaMP7b (middle) or with jGCaMP7b-X_C_ (right). At -10 mV, Ca^2+^ current traces (red, with scale bars for current amplitudes) and Ba^2+^ current traces (gray, rescaled) are shown. *SCa* and *ICa* (quantified by the equations shown in the first column) are the indices of calcium dependent inactivation and voltage gated activation, respectively. Standard error of the mean (S.E.M) and two-tailed unpaired Student’s *t*-test (**B**) or one-way ANOVA followed by Bonferroni for post hoc tests (**C-E**) (criteria of significance: \**p*<0.05, \*\*\**p*<0.001) were applied.

### Acute sensory responses monitored by viral expression of GCaMP-X *in vivo*

GECIs including GCaMP have been widely applied to monitor neuronal responses to various stimuli. Due to the cytotoxicity known from the very early versions of GCaMP, *in vivo* imaging experiments are normally required to conduct within the time window. In practice, an optimal time window (OTW) is about 3-8 weeks post injection for GCaMP-infected neurons of live mice (Chen et al., 2013; Resendez et al., 2016), in order to achieve substantial levels of GCaMP expression/fluorescence but not too high levels prone to side effects. Here we investigated into the Ca^2+^ dynamics under whisker stimulation within or beyond OTW by applying the adeno-associated viruses (AAV) of GCaMP6m or GCaMP6m-X_C_ with the neuro-specific *Syn* promoter to S1 primary somatosensory cortex in mouse brain (***Figure 2A***). GCaMP6m was aggregated in the nucleus of a fraction of the neurons, especially beyond OTW, consistent with the progressive nuclear accumulation previously observed *in vivo* and *in vitro* (Chen et al., 2013; Yang et al., 2018). 4 weeks post injection, the nucleus-filled neurons were around 16% among the total, and then rose up to 33% when approaching 10 weeks post injection (***Figure 2B***). In direct contrast, nuclear accumulation was barely noticeable from GCaMP6m-X_C_ even beyond OTW (up to 11 weeks post injection). In the earlier study (Chen et al., 2013), the responses to visual stimuli from neurons expressing nucleus-filled GCaMP6 were impaired after several months post injection; meanwhile, it was reported that during the initial phase (1-2 month post injection) nuclear GCaMP would not perturb the proper physiology of neurons. However, our data demonstrated that the sensory neurons here, when filled with nuclear GCaMP, were less responsive than neurons expressing GCaMP-X_C_ no matter within or beyond OTW (***Figure 2A)***, as quantified by the success rate in response to repetitive whisker stimuli (***Figure 2C***). Moreover, for neurons expressing GCaMP mostly in the cytosol (4 weeks post injection), the average amplitude of Ca^2+^ responses (*ΔF/F0*) was significantly lower (*p*<0.05) (***Figure 2C***), in agreement with the *in vitro* results that cytosolic GCaMP would bind apoCaM targets such as CaV1 channels thus perturbing crucial signals and cell functions (Yang et al., 2018). Regarding the average amplitude of sensory responses beyond OTW, no significant difference was found between cytosolic GCaMP and GCaMP-X_C_ which may involve certain compensatory effects as being postulated. However, the overall performance of GCaMP is still inferior to GCaMP-X_C_ considering its significantly lower success rate.

**Figure 2.**
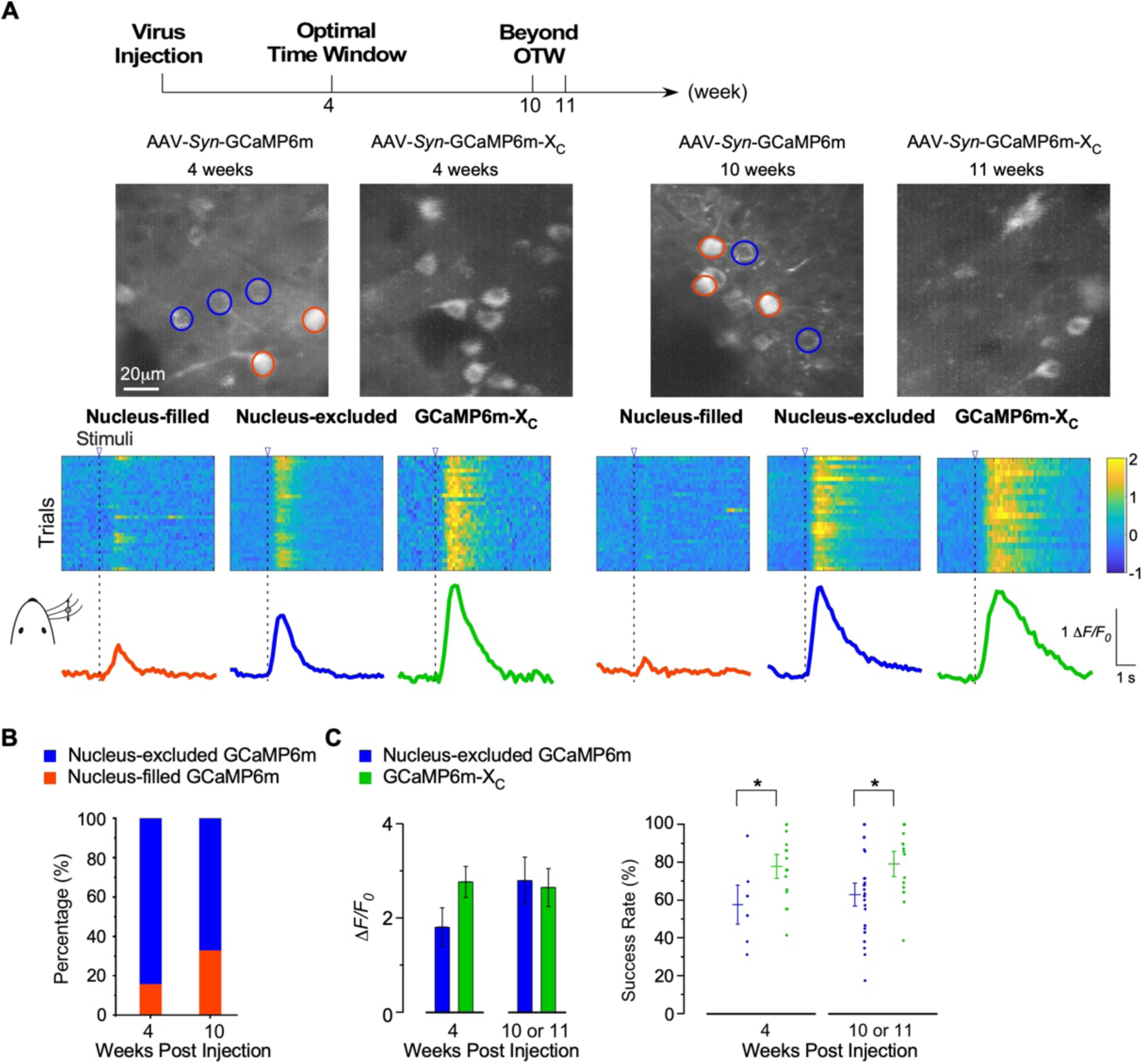
Ca^2+^ imaging of sensory-evoked responses in GCaMP-X versus GCaMP virus-infected neurons *in vivo*. (**A**) Ca^2+^ dynamics of GCaMP6m and GCaMP6m-X_C_ in S1 primary somatosensory cortex under whisker stimulation by *in vivo* two-photon Ca^2+^ imaging. 4 weeks and 10-11 weeks post injection are considered as optimal time window (OTW) or beyond OTW, respectively. The orange and blue circles in the representative two-photon images indicate nucleus-filled and nucleus-excluded GCaMP6m, respectively. The colored scale bar indicates the fluorescence intensity of Ca^2+^ probes. Multiple trials of the same neuron are shown for nucleus-filled GCaMP6m, nucleus-excluded GCaMP6m, or GCaMP6m-X_C_. The averaged Ca^2+^ responses are shown at the bottom. (**B**) Percentage of nucleus-excluded versus nucleus-filled neurons expressing GCaMP6m, within or beyond OTW. (**C**) Responses to repetitive whisker stimuli were evaluated and compared for neurons expressing GCaMP6m (nucleus-excluded) versus GCaMP6m-X_C_. The average amplitude (*ΔF*/*F0*) (left) and success rate (right) of the trials. Standard error of the mean (S.E.M) and Student’s *t*-test (criteria of significance: * *p*<0.05) were calculated.

Therefore, our observations extend the previous notions mostly obtained from *in vitro* data onto *in vivo* GCaMP imaging. Neurons still suffer from the side-effects if simply relying on OTW and/or the index of nuclear-cytosolic distribution. Overall, GCaMP-X outperforms GCaMP in imaging sensory-evoked Ca^2+^ dynamics, which stands out with its high neuron-compatibility suitable to long-term Ca^2+^ imaging.

### Long-term monitoring of Ca^2+^ oscillations by GCaMP-X_C_ *in vitro*

Before proceeding further with the actual *in vivo* imaging with GCaMP-X, we decided to conduct in-depth examinations on the long-term performance of GCaMP-X under *in vitro* conditions. Of note, slow Ca^2+^ oscillations have been observed from a variety of excitable or non-excitable cells (Uhlen and Fritz, 2010). In neurons, oscillatory Ca^2+^ signals can increase the efficiency and specificity of gene expression (Dolmetsch et al., 1998; Li et al., 1998), thus playing important role in neurogenesis (Kamijo et al., 2018; Toth et al., 2016). Meanwhile, spontaneous electrical activities are initiated at the early stage of neural development, and subsequently become more synchronized (Luhmann et al., 2016; Spitzer, 2006). We hypothesized that GCaMP-X versus GCaMP neurons would make a clear difference in their chronic profiles of Ca^2+^ oscillations, as the potential mechanistic linkage between channel dysregulation and neuronal impairment, both evidenced from GCaMP-expressing neurons. Meanwhile, neuron-compatible GCaMP-X is expected to provide new insights into the roles of spontaneous Ca^2+^ oscillations in neuronal development such as neurite morphogenesis (Gomez and Zheng, 2006; Rosenberg and Spitzer, 2011).

To monitor such Ca^2+^ dynamics *in vitro*, the adeno-associated viruses (AAV) carrying GCaMP6 or GCaMP6-X_C_ (for *in-vitro* use) were applied to cortical neurons of neonatal mice (DIV 0, 0 day *in vitro*) which were cultured for DIV 28 or longer (***Figure 3A, B***). Fluctuations of Ca^2+^ activities were perceivable starting from the first week (DIV 3 and DIV 6) with GCaMP6-X_C_, in the pattern of high-frequency, low-amplitude and unsynchronized signals (***Figure 3—video supplement 1***). On DIV 10, the oscillation frequency decreased while the amplitude was increased with an enhanced level of synchronization (***Figure 3—video supplement 2***). On DIV 28, Ca^2+^ oscillations of individual or sub-grouped neurons became broadly synchronized featuring robust spikes and slow frequency (***Figure 3—video supplement 3***), indicative of the formation of neural circuitry. In contrast, Ca^2+^ signals were severely distorted in GCaMP6-infected DIV-10 neurons. Despite that the performance of GCaMP6 in the first week resembled GCaMP6-X_C_ (***Figure 3—video supplement 4***), longer expression time of GCaMP6m resulted in altered patterns of Ca^2+^ oscillations (***Figure 3—video supplement 5, 6***). One major abnormity was the substantial reduction in oscillatory activities of GCaMP6-expressing neurons, which was manifested after DIV 10 by much longer intervals between two adjacent peaks and much smaller amplitudes in average. Occasionally, abnormal Ca^2+^ spikes with ultralong lasting duration could be observed on DIV 17 (***Figure 3B***). At the end of one month (DIV 28), there was barely any recognizable Ca^2+^ oscillation and broken neurites were often evidenced (***Figure 3—video supplement 6***). We then further analyzed oscillatory Ca^2+^ signals by the frequency and other key indices across the timespan from DIV 3 up to DIV 28. Statistical results demonstrated that the frequency of Ca^2+^ fluctuation with GCaMP6-X_C_ was around 150 mHz during the first week, then gradually declined to the plateau around 20 mHz (***Figure 3C***). Regarding the peak amplitude, a persistent rising trend was evidenced from GCaMP6m-X_C_ across the full term (***Figure 3D***). In contrast, both the frequency and the amplitude of Ca^2+^ oscillations acquired by GCaMP6m were drastically changed after DIV 17, and then even more deteriorated later in that the oscillation was less and less recognizable and eventually halted on DIV 28 (***Figure 3C, D***). Synchronization is one major hallmark of autonomous Ca^2+^ oscillations, which was evaluated by the index of the mean of correlation coefficient. As demonstrated by the temporal profiles of correlation coefficients, the comparison between GCaMP6m versus GCaMP6-X_C_ unveiled a similar pattern to that of the amplitude, which depicted the crucial phase around DIV 17-20 with a clear tendency of reduction (***Figure 3E***). Likewise, the full width at half maximum (FWHM), another index of Ca^2+^ waveforms, was aberrantly wider for GCaMP6 than GCaMP6-X_C_, becoming noticeable on DIV 10, and much more pronounced (10-fold) later on (***Figure 3F***). Collectively from these indices, GCaMP indeed caused detrimental side-effects on the neurons along with culturing time or developmental stages; in contrast, GCaMP6-X_C_ has overcome nearly all the above negative effects, emerging as a neuron-compatible tool for chronic Ca^2+^ imaging. Also, Fast Fourier Transformation (FFT) was applied to the Ca^2+^ waveforms acquired by GCaMP6-X_C_ (***Figure 3G*** and ***Figure 3—figure supplement 1***). The distribution of frequency components started to change during DIV 10-17, when slow Ca^2+^ oscillations of 10-100 mHz appeared to be the dominant form (***Figure 3H***). With independent preparations of neurons, additional sets of long-term imaging experiments with GCaMP6-X_C_ up to DIV 42 (***Figure 3—figure supplement 2***), or with the newer probes of jGCaMP7 and jGCaMP7-X_C_ and an extended timeframe of DIV 35 (***Figure 3—figure supplement 3***), led to consistent results supporting that GCaMP-X outperforms GCaMP in chronic Ca^2+^ imaging. Here, the same set of neurons are chronically monitored with GCaMP-X across the full term (one month or longer), in order to establish the temporal profiles of the major characteristics for autonomous Ca^2+^ oscillations.

**Figure 3.**
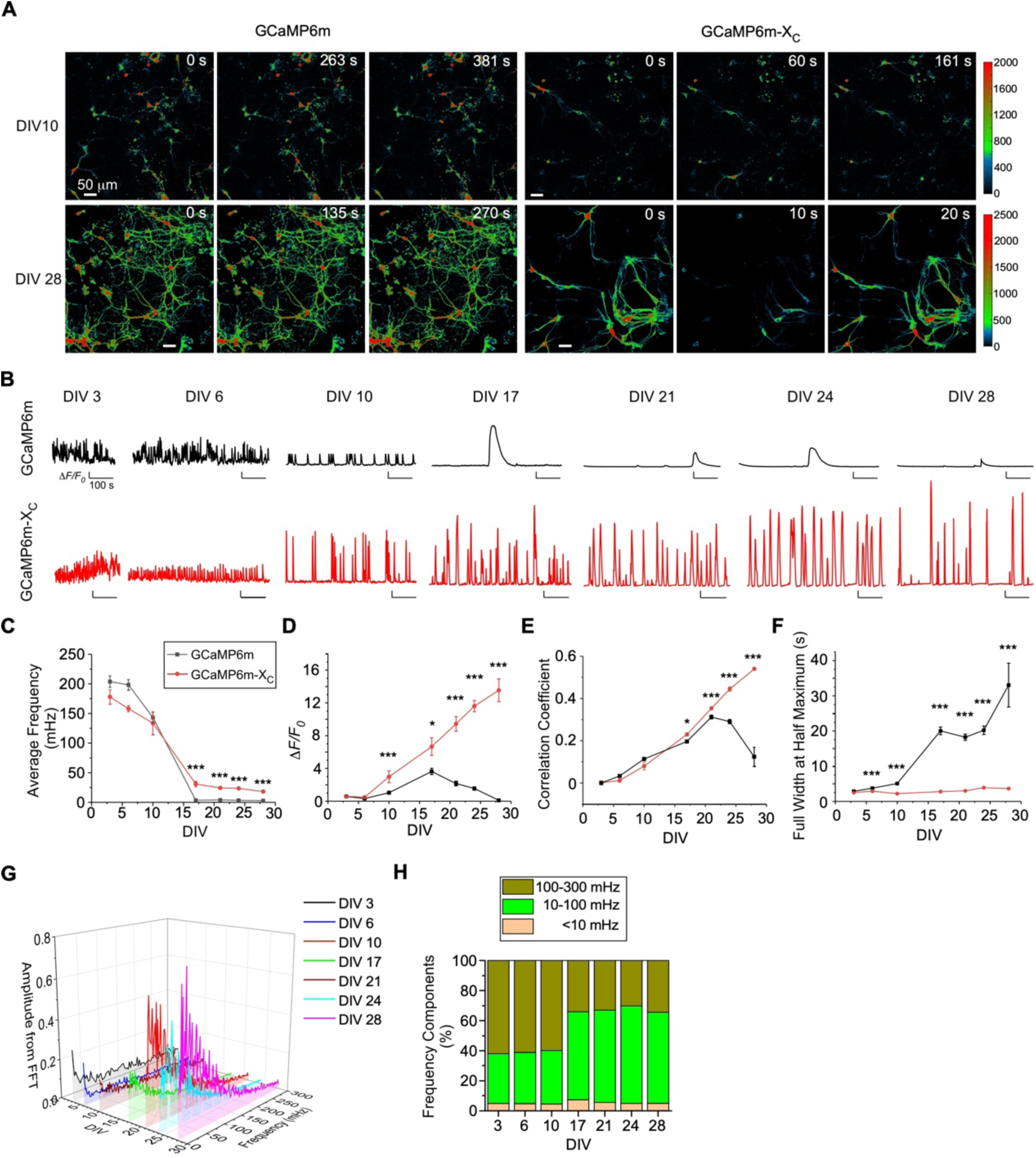
Chronic Ca^2+^ fluorescence imaging for autonomous Ca^2+^ oscillations in cultured cortical neurons. (**A**) Time-lapse images of cultured cortical neurons infected with AAV-*Syn*-GCaMP6m or AAV-*Syn*-GCaMP6m-X_C_. Spontaneous Ca^2+^ activities by the two probes are shown for DIV-10 and DIV28. See Figure 3***—video supplement 2, 3, 5*** and ***6*** for details. Ca^2+^ signals (color-coded) were monitored by a confocal microscope with a live-cell imaging chamber to maintain the conditions for the cell culture (37°C, 5% CO2, 97-100% humidity) at different time-points as marked. (**B**) Representative traces of Ca^2+^ activities in cultured cortical neurons expressing GCaMP6m (upper) or GCaMP-X_C_ (lower) from DIV 3 to DIV 28. (**C-F**) Temporal profiles of key indices measured from spontaneous Ca^2+^. The average frequency (10^-3^ Hz or mHz, **C**) and peak amplitude (*ΔF/F0*, **D**); synchrony (quantified by the mean of correlation coefficient per view, **E**), and full width at half maximum (FWHM, **F**). (**G**) Power spectral analyses by FFT (Fast Fourier Transformation) for Ca^2+^ traces of cortical neurons from DIV 3 to DIV 28. (**H**) Frequency components in percentage. By integrating the absolute amplitudes over each frequency band (**G**), three major bands are shown: <10 mHz (ultra-slow), 10-100 mHz (slow) and 100-300 mHz (fast), where 10-100 mHz is considered as the major component. Standard error of the mean (S.E.M) and the Student’s *t*-test (**C-F**) (two-tailed unpaired with criteria of significance: * *p*<0.05 and \*\*\**p*<0.001) were calculated when applicable.

### Close correlations between autonomous Ca^2+^ oscillations and neuronal development *in vitro*

Spontaneous Ca^2+^ oscillations, the slow periodic Ca^2+^ waveforms in particular, are tightly coupled with neuronal development including neuritogenesis (Kamijo et al., 2018; Toth et al., 2016). GCaMP-X promises unprecedented opportunities for concurrent imaging of both neuronal functionalities and morphogenesis during the long term of development. Such chronic Ca^2+^ imaging is infeasible if using other approaches, e.g., conventional GCaMP or Ca^2+^ dyes, both would bring side-effects to neurons (Smith et al., 2018; Yang et al., 2018). By taking advantage of GCaMP-X, we here aimed at the inter-relationship between cellular Ca^2+^ and neuronal development.

Indistinguishable from control neurons infected with GFP viruses (***Figure 4—figure supplement 1***), neurons expressing GCaMP6m-X_C_ followed the typical development process of neonatal neurons, including neurite elongation/arborization and soma enlargement (***Figure 4A***). In contrast, these developmental processes were severely impaired by virally-delivered GCaMP6m, especially after DIV 14 onwards, when nuclear accumulation and neurite shortening became evident. Depicted by DIV-28 neurons with neurite tracings, GCaMP6m caused significant damages on neurite outgrowth, and to the extreme, discernable death of neurons, in contrast to GCaMP6m-X_C_ which caused no apparent perturbation. In addition, the temporal profiles across the full time-course were shown for both GCaMP6m (***Figure 4B***) and GCaMP6m-X_C_ (***Figure 4C***) by the major indices of total neurite length and soma size. At the early phase (before DIV 17), no significant difference between the two groups of GCaMP6m versus GCaMP6m-X_C_ could be observed; however, toward DIV 28 of GCaMP-expressing neurons, the soma size could be as small as ∼200 μm^2^ in contrast to the neurons expressing GFP or GCaMP6m-X_C_ (∼300 μm^2^) (**Figure 4—figure supplement 1**). Likewise, the total neurite length of GCaMP-expressing neurons rapidly declined, whereas GCaMP-X expressing neurons went through an initial phase (∼ 2 weeks) of rapid outgrowth before entering into the plateau phase, indicated by the trending curve. Similar to neuritogenesis, the soma size also exhibited a temporal profile of upward-plateau pattern (***Figure 4C***). Combining the data and analyses from both developmental and functional perspectives (***Figure 3*** and ***Figure 4***), we could speculate the potential correlations between neuronal growth and spontaneous Ca^2+^ activities (***Figure 4—figure supplement 2***). Functionally, the amplitude and the frequency appeared to be either ascending (amplitude) or descending (frequency) along with the developmental stage (DIV) (***Figure 4D***). Roughly, the oscillation amplitude linearly (*R^2^*=0.84) correlated with the neurite length in total (***Figure 4—figure supplement 2A***). In direct contrast, the oscillation frequency and neurite outgrowth (total length) were inversely correlated (*R^2^*=0.99) (***Figure 4—figure supplement 2B***). Resembling the amplitude, the level of synchrony indicative of circuitry formation was positively correlated with the total neurite length (*R^2^*=0.86) (***Figure 4— figure supplement 2C***). All these tight correlations support the notion that spontaneous Ca^2+^ activities including its mature form of autonomous Ca^2+^ oscillations may underpin the neuritogenesis (Estrada et al., 2006; Gomez and Zheng, 2006; Kamijo et al., 2018). To be more specific, based on our earlier spectral analyses (***Figure 3G, H***), the major component of 10-100 mHz (rather than the other components) stood out as the major driving force of neuritogenesis.

**Figure 4.**
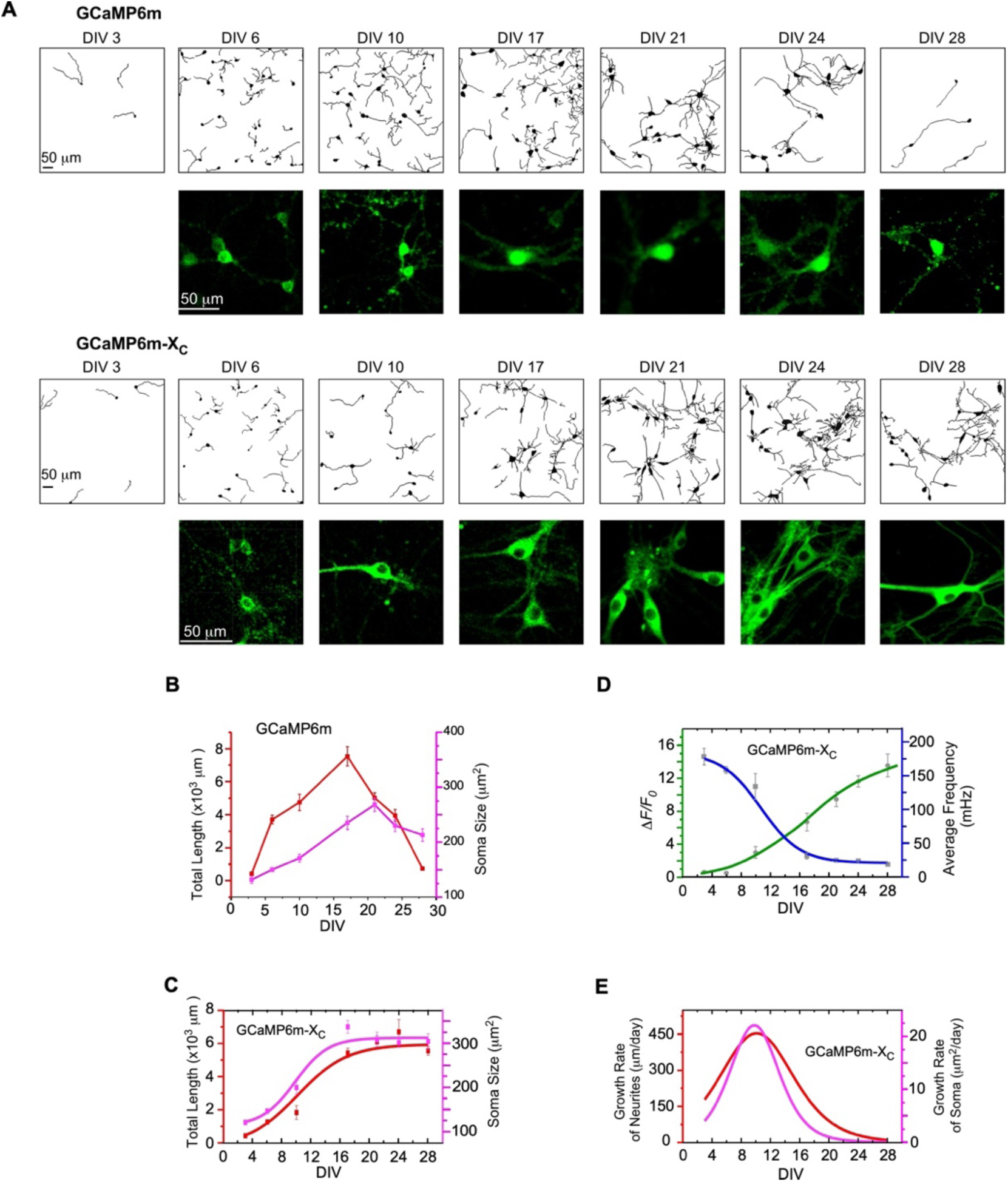
The correlations between neuronal development and Ca^2+^ oscillations. (**A**) Time-lapse images with neurite tracing for cultured cortical neurons expressing GCaMP6m (upper two rows) or GCaMP6m-X_C_ (lower two rows). Enlarged fluorescence images are to show subcellular distributions of the probes, indicative of the nuclear accumulation of GCaMP6m versus GCaMP6m-X_C_. (**B**) Temporal profiles of total neurite length per view (red) or soma size per neuron (purple) for neurons expressing GCaMP6m. (**C**) Temporal profiles of total neurite length per view (red) or soma size per neuron (purple) for neurons expressing GCaMP6m-X_C_. (**D**) The temporal profiles of the average frequency of Ca^2+^ oscillations (blue) and the peak amplitude (*ΔF/F0*, green), adopted from Figure 3C***, D***. (**E**) Temporal profiles of the growth rates (μM/day) of neurite length (red) or soma size (pink), respectively. Standard error of the mean (S.E.M) was calculated when applicable.

Previous studies mainly rely on measuring the transient Ca^2+^ and growth rate of neurites within a brief period of time (Ito et al., 2010; Mukai et al., 2010; Rosenberg and Spitzer, 2011; Van et al., 2004). However, the overall neurite outgrowth across the developmental phases may help elucidate the roles of Ca^2+^ in neuritogenesis, which has been largely unavailable due to lack of chronic Ca^2+^ imaging. In fact, contradictory observations have been reported regarding how Ca^2+^ actually regulates the rates of neuronal growth. Here, GCaMP-X imaging enabled us to calculate the first derivative of the time-dependent neurite/soma growth as the growth rate of neurite or soma (***Figure 4E***). The resulted bell-shaped curves unveiled that the growth rates could be in distinct relationships with Ca^2+^ oscillations, depending on the particular stage of development, which reached the peak rates around DIV 10-12 for both neurite length and soma size. Therefore, it seems there exists an optimal combination of amplitude and frequency, by which Ca^2+^ oscillations could drive the growth of neurites and soma at its maximum rate. In general, there might not be a simple relationship between Ca^2+^ oscillation characteristics and neuronal development. Without concurrent information on both neuritogenesis and Ca^2+^ signals, it could be rather complicated or even controversial if the correlations are attempted between growth rates and various combinations of amplitude and frequency within an acute or transient timeframe (Gomez and Zheng, 2006). Here, the roles of slow Ca^2+^ oscillations in neuritogenesis started to unveil, in the light of chronic GCaMP-X imaging.

### Chronic imaging of spontaneous Ca^2+^ activities *in vivo*

Similar to cultured neurons, spontaneous Ca^2+^ activities *in vivo* are also correlated to gene transcription and expression at the cellular and circuity levels (Laviv et al., 2020; Takahashi et al., 2016). By applying AAV viruses carrying GCaMP6m or GCaMP6m-X_C_ to the mouse brain, we characterized spontaneous Ca^2+^ activities in S1 primary somatosensory cortex (***Figure 5A, B*** and ***Figure 5—video supplement 1-4***). Two checkpoints were set at 4-weeks post virus injection (within OTW) and at 8- or 11-weeks (prolonged expression time beyond OTW), respectively. Similar to whisker deflection-response experiments in ***Figure 2***, the nucleus-filled neurons exhibited noticeable abnormalities in spontaneous Ca^2+^ activities even within OTW; and the damages were much exacerbated beyond OTW. The frequency, amplitude, and SNR (signal-to-noise ratio) were significantly lower, accompanied by aberrantly wider FWHM (full width at half maximum) and slower on/off rates (***Figure 5C***). In contrast, neurons expressing GCaMP6m-X_C_ maintained robust and stable spontaneous Ca^2+^ activities with key characteristics of normal values across the full term of experiments (up to 11 weeks post virus injection). Notably, GCaMP6m-X_C_ significantly improved the SNR of Ca^2+^ signals *in vivo*, manifested by 2-fold SNR of nucleus-excluded GCaMP6m and several times higher than nucleus-filled GCaMP6m (***Figure 5C***).

**Figure 5.**
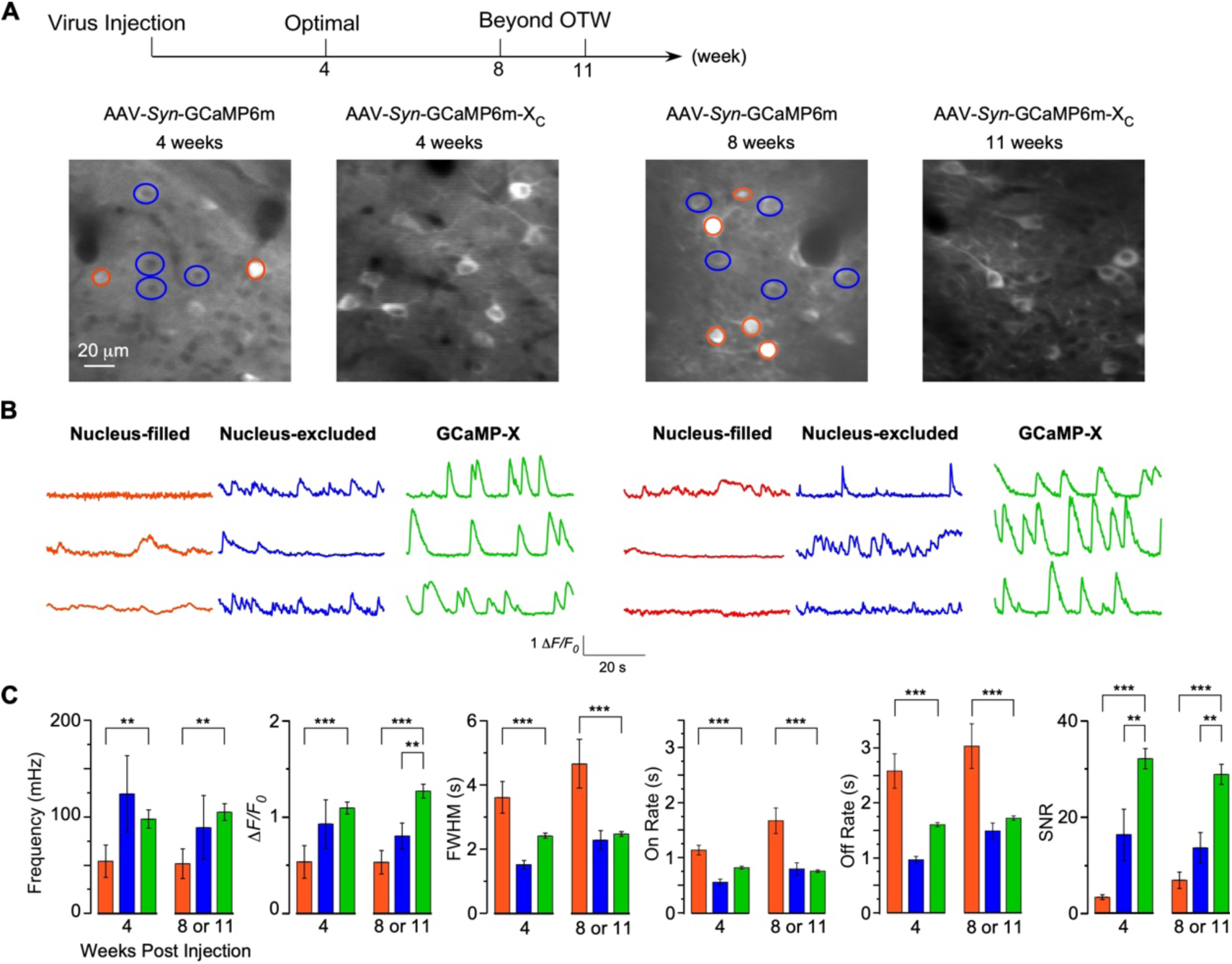
Chronic Ca^2+^ imaging for spontaneous Ca^2+^ activities *in vivo*. (**A, B**) *In vivo* two-photon fluorescence images (**A**) and spontaneous nuclear Ca^2+^ activities (**B**) of virus-infected neurons in S1 primary somatosensory cortex. Neurons expressing GCaMP6m-X_N_ or GCaMP6m (nucleus-filled and nucleus-excluded) within OTW (optimal time window) and beyond OTW are compared. (**C**) Key parameters of spontaneous Ca^2+^ activities in (**B**). Standard error of the mean (S.E.M) and one-way ANOVA followed by Bonferroni for post hoc tests (criteria of significance: ** *p*<0.01, and \*\*\**p*<0.001) were calculated when applicable.

Inspired by chronic GCaMP-X imaging *in vitro*, we speculated a similar correlation between Ca^2+^ activities and neuronal development which may underlie GCaMP impairments *in vivo*. Firstly, titrations of GCaMP and GCaMP-X viruses were applied to characterize the dose-dependent damages of neurons, for which the concentrations were at 1x10^11^, 5x10^11^ and 1x10^12^ v.g./ml for AAV-*Syn*-GCaMP6m, and 1x10^12^ v.g./ml for AAV-*Syn*-GCaMP6m-X_C_, respectively. 30 nl viruses at the above concentrations were microinjected into different brain regions of the same mouse and then after three weeks the expression levels in brain slices were examined (***Figure 6A***). Low-concentration injection of virus at 1x10^11^ v.g./ml exhibited extremely sparse expression of GCaMP6m and yielded a low cell count. Correspondingly, the fluorescence signals were difficult to distinguish from the background, i.e., the ultra-low contrast. The virus concentration, when increased to 5x10^11^ v.g./ml, resulted in a relatively larger number of healthy-looking cells expressing GCaMP6m. But the low image contrast still affected proper detection of Ca^2+^ signals due to enhanced neuropil fluorescence. High-expression levels of GCaMP (at the virus concentration of 1x10^12^ v.g./ml) significantly enhanced the fluorescence image contrast (SNR) and greatly increased the numbers of GCaMP-positive cells. However, the majority of neurons exhibited severe nuclear accumulation, which would subsequently lead to aberrant Ca^2+^ dynamics and cell death. In contrast, high-dose injection of GCaMP6m-X_C_ virus at 1x10^12^ v.g./ml was beneficial for image contrast and the number of positive and healthy cells; meanwhile the N/C ratio remained ultra-low as expected (***Figure 6B***). Next, we injected high doses of GCaMP6m (1x10^12^ v.g./ml) and ultra-high doses of GCaMP6m-X_C_ (1x10^13^ v.g./ml, 10-fold higher) viruses to examine the temporal profile of the damages (***Figure 6C***). At the checkpoints of 17-, 55-, 70- and 92-days post injection, neurons expressing GCaMP6m-X_C_ were compared with GCaMP6m. Confocal microscopy with brain slices revealed that the percentage of infected neurons and the expression level of GCaMP6m-X_C_ were close to their peaks on 17-days, suggesting that the ultra-high dose could expedite GCaMP6m-X_C_ expression to reach the high level. Most importantly, long-term, high-level expression of GCaMP6m-X_C_ up to 92-days did not induce nuclear accumulation, whereas GCaMP6m at relatively lower concentration (1x10^12^ v.g./ml) already caused severe nuclear accumulation evidenced from 17-days to 92-days (***Figure 6D***). Meanwhile, the soma size of GCaMP6m-infected neurons was significantly smaller than that of GCaMP6m-X_C_ from 55-days to 92-days, presumably due to impaired Ca^2+^ signaling (including spontaneous Ca^2+^ activities) in these neurons. In summary, GCaMP-X_C_ has the high compatibility with neurons as desired by chronic Ca^2+^ imaging, validated by the full set of tests of both *in vitro* and *in vivo* and of both acute sensory responses and spontaneous Ca^2+^ fluctuations.

**Figure 6.**
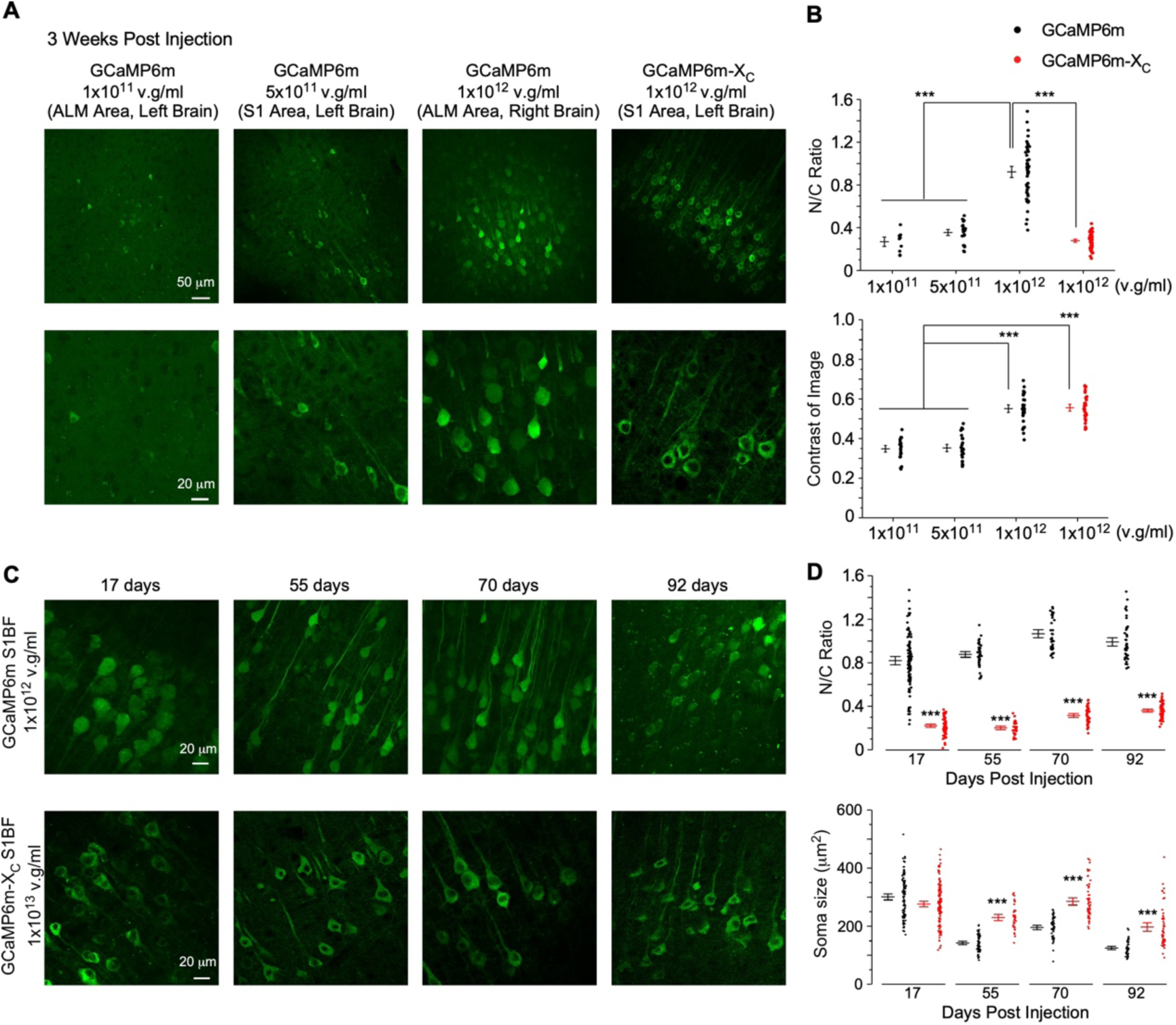
Chronic evaluation of neuronal growth *in vivo*: GCaMP-X versus GCaMP. (**A**) Confocal images of the brain slice from mice injected with different doses of GCaMP6m or GCaMP6m-X_C_ viruses (30 nl). Brain slice were dissected 3 weeks after virus injection. (**B**) N/C ratio (upper) and contrast of images (lower). (**C**) Confocal images of brain slices expressing GCaMP6m or GCaMP6m-X_C_ acquired at different timepoints up to 92 days post virus injection. AAV-*Syn*-GCaMP6m (1.0x10^12^ v.g./ml, 60 nl) or AAV-*Syn*-GCaMP6m-X_C_ (1.0x10^13^ v.g./ml, 60 nl) was microinjected into left or right S1BF (the barrel field of the primary somatosensory cortex) of the same mouse, respectively. Brain slices of injected mice were dissected and compared. (**D**) Summary of the N/C ratio (left) and soma size (right) for the neurons expressing GCaMP6m or GCaMP6m-X_C_. Standard error of the mean (S.E.M) and one-way ANOVA followed by Bonferroni for post hoc tests (criteria of significance: ** *p*<0.01, and \*\*\**p*<0.001) were calculated when applicable.

### GCaMP side-effects in transgenic mice unveiled by cultured cortical neurons

Although the drawbacks of GCaMP were noticed at the very beginning and then mechanistically resolved later on, GCaMP transgenic mice have been considered to be relatively safe in comparison with viral delivery of GCaMP. Nevertheless, recent studies reported that some transgenic mouse lines, such as Ai93 and Ai148, suffered from epileptiform activities (Daigle et al., 2018; Steinmetz et al., 2017), often in association with long-term and/or high-level expression of GCaMP in neurons. Based on our earlier study on transient and viral overexpression, we suspected that the mechanisms of side-effects should be applicable to transgenic expression of GCaMP. In this work, brain slice or culture neurons from transgenic mice were examined from both functional and morphological aspects.

Ai148 is a widely-used transgenic line, for which TIGRE2.0 has been utilized for GCaMP6f to enhance its expression level, such that the potential damages by GCaMP are expected to be more pronounced (Daigle et al., 2018). We examined the 6-month old Rasgrf2-2A-dCre;Ai148D mice (with TMP-inducible expression of GCaMP6f), by comparing GCaMP-expressing neurons with GFP-infected control neurons from the layer II-III cortex (***Figure 7A***). Nuclear accumulation of GCaMP was readily discernible, although it was less severe than high-dose viral expression of GCaMP6m (***Figure 6***). Consistent with the ectopic distribution of GCaMP, Ai148 neurons exhibited significant reduction in soma size relative to GFP control neurons (***Figure 7B***), altogether supporting that Ai148 mice still suffer from intrinsic GCaMP side-effects.

**Figure 7.**
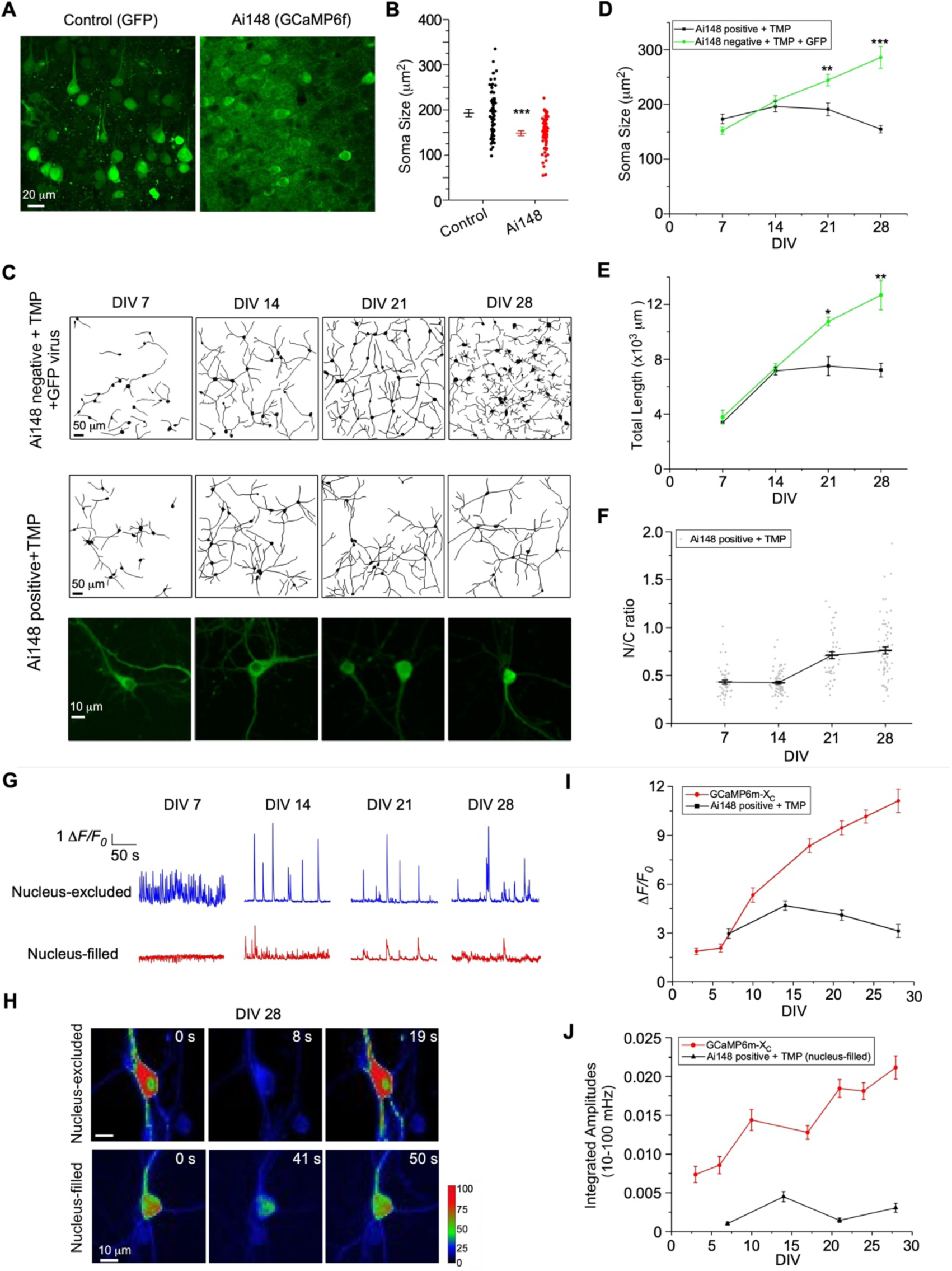
Chronic imaging for cortical neurons from GCaMP transgenic mice. (**A**) Brain slice images of cortex layer II-III from 6-month old mice injected with GFP virus (control, left), or from 6-month old GCaMP6f-positive transgenic mice (Rasgrf2-2A-dCre x Ai148, right). GCaMP6f was examined ∼3 weeks after induction of expression by intraperitoneal injection of TMP. (**B**) The soma size of layer II-III neurons of GFP control mice versus GCaMP6f transgenic mice. (**C**) Neuronal growth of cultured cortical neurons from Ai 148 mice. Neurite tracing for GCaMP-negative neurons (infected by GFP virus) versus GCaMP-positive neurons, added with 10 μM TMP for expression induction. Confocal images of individual neurons are zoomed in to illustrate subcellular distributions of GCaMP (bottom row). (**D-F**) Temporal profiles of total neurite length (**D**), soma size (**E**), or N/C ratio (**F**) for GCaMP-expressing Ai148 neurons, in comparison with GCaMP-negative GFP-infected neurons, both shown in (**C**). (**G**) Spontaneous Ca^2+^ activities of GCaMP-positive neurons in either nuclear-excluded or nuclear-filled subgroups (with the criteria of N/C ratio<0.6 or >0.8). (**H**) Exemplars of single-neuron Ca^2+^ images (color-coded) shown for DIV-28 neurons (nucleus-excluded versus nucleus-filled). (**I** and **J**) Statistical profiles of the peak amplitude (in Δ*F/F0*, **I**) and the integrated amplitudes (within the frequency band of 10-100 mHz, **J**) for Ai148 GCaMP-positive neurons, compared with GCaMP-X_C_. Standard error of the mean (S.E.M) and Student’s *t*-test (**B, D** and **E**) (two-tailed unpaired with criteria of significance: * *p*<0.05, ** *p*<0.01, *** *p*<0.001) were calculated when applicable.

Next, dissected from newborn Rasgrf2-2A-dCre;Ai148 mice, cortical neurons were cultured, and subsequently 10 μM TMP was added to induce GCaMP expression. Similar to viral delivery, transgenic GCaMP6f neurons also perturbed neuronal development, especially for nucleus-filled neurons. Neurite tracings suggested that the complexity and length of neurites were significantly reduced in transgenic GCaMP neurons as compared to GFP control neurons starting from DIV 21 (***Figure 7C***). Temporal profiles of neuronal development revealed that the growth of soma size and neurite length significantly slowed down or even halted on DIV 14, in comparison with the control neurons (GFP virus-infected and TMP treated) (***Figure 7D, E***). Consistently, N/C ratio of GCaMP indicative of nuclear accumulation was gradually increased along with the culturing time up to one month (***Figure 7F***). Functionally, Ca^2+^ waveforms of much lower amplitude were imaged from Ai148 neurons expressing nucleus-filled GCaMP across the full month than the nucleus-excluded subgroup, further supporting the notion learned from GCaMP plasmids and viruses that the side-effects would be exacerbated by nuclear GCaMP (***Figure 7G***, **Figure 7—video supplement 1** and **Figure 7—video supplement 2**). Similar results were obtained from analyzing the peak amplitude and integrated frequency of Ca^2+^ oscillations by comparing nucleus-filled versus nucleus-excluded subgroups of neurons on DIV 14 or later (***Figure 7H-J***). Another trial of neurite tracing and Ca^2+^ imaging was performed with TMP-inducible expression of GCaMP, which confirmed the effects and analyses described above (**Figure 7—figure supplement 1)**.

In addition to chemical-inducible expression of GCaMP, newborn Emx1-Cre;Ai148 mice were deployed to constitutively express GCaMP6f (**Figure 7—figure supplement 2**). Persistent GCaMP6f expression is expected to cause similar damages to cortical neurons. Indeed, GCaMP6f started to accumulate in the nucleus at early stage (0.57±0.03 on DIV 4), but did not exacerbate as checked on DIV 28 (0.59±0.02). Even so, the neuronal damages were clearly evidenced when compared with those by GCaMP6m-X_C_ viruses. Meanwhile, the major characteristics of spontaneous Ca^2+^ oscillation in transgenic neurons were also significantly altered, resulting in relatively lower frequency, less synchronization, smaller amplitude, and abnormally increased FWHM.

In summary, the major findings by virally-transfected neurons are applicable to two different lines of transgenic mice, supported by morphological and/or functional analyses from both brain slices of adult mice and long-term culturing of transgenic neurons: cortical neurons of transgenic mice are also perturbed by GCaMP, causing neuronal damages similar to those by GCaMP viruses.

### Applications of nucleus-localized GCaMP-X_N_ *in vitro* and *in vivo*

Highly localized Ca^2+^ signals in different cellular organelles play subtle and specific roles in diverse processes, e.g., nuclear Ca^2+^ directly regulates gene expression and neuronal development (Bading, 2013). Conventional tools to monitor nuclear Ca^2+^ signal is based on nuclear-targeted GECIs, by tagging the nuclear localization sequence, i.e., GCaMP-NLS ( (Kim et al., 2014). However, such nuclear probes would cause severe damages to mammalian neurons and distort Ca^2+^ signals (Yang et al., 2018). GCaMP-X_N_ or the nuclear version of GCaMP-X has been proposed as a more faithful tool to reflect authentic Ca^2+^ dynamics in the nuclei (Yang et al., 2018). Here, we applied AAV-*Syn*-GCaMP6m-X_N_ viruses to cultured cortical neurons for multi-week examinations of nuclear Ca^2+^ (***Figure 8A***). Similar to cytosolic GCaMP6m-X_C_ (***Figure 3***), the nuclear Ca^2+^ waveforms at the early stage of development (DIV 7) were unsynchronized, of high frequency and low-amplitude (***Figure 8B***). On DIV-28, GCaMP-X_N_ demonstrated that spontaneous Ca^2+^ activities transformed into highly-synchronized oscillations of slower rate and higher amplitude. In contrast, cortical neurons with nucleus-filled GCaMP6m barely exhibited any oscillatory activities (***Figure 8C-E***).

**Figure 8.**
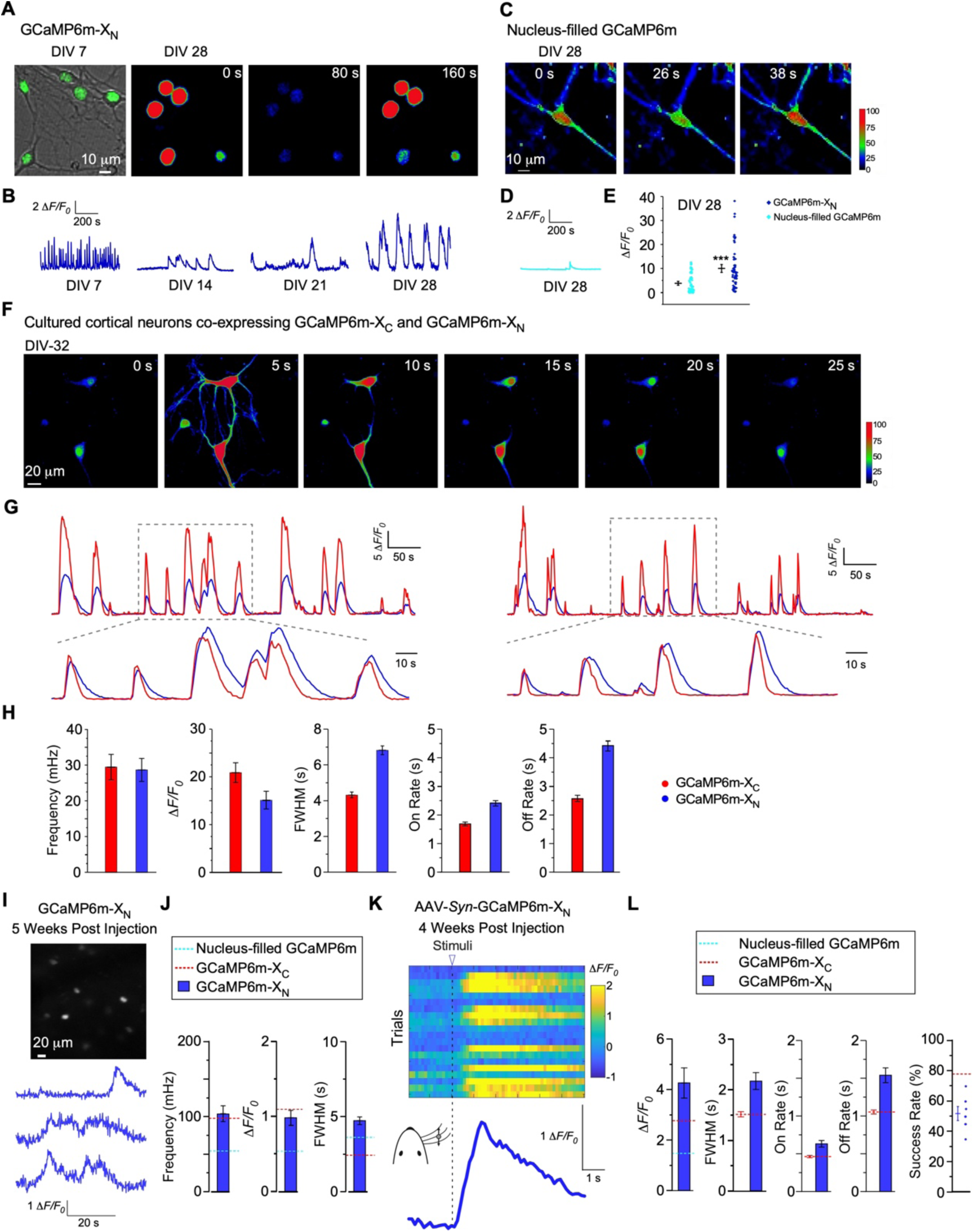
Chronic GCaMP-X_N_ imaging of nuclear Ca^2+^ *in vitro* and *in vivo*. (**A** and **B**) Wide- field image superimposed with GCaMP-X_N_ fluorescence image (**A**, left), color-coded Ca^2+^ fluorescence images of one oscillation cycle (**A**, right), and spontaneous Ca^2+^ waveforms (**B**) from cultured cortical neurons infected with AAV-*Syn*-GCaMP6m-X_N_. (**C** and **D**) For DIV-28 neurons filled with AAV-*Syn*-GCaMP6m in the nucleus, color-coded Ca^2+^ fluorescence images (**C**) and irregular Ca^2+^ waveforms (**D**). (**E**) The amplitudes of spontaneous Ca^2+^ waveforms by GCaMP-X_N_ versus nuclear GCaMP. (**F**) Confocal fluorescence images of cultured cortical neurons infected with both GCaMP6m-X_C_ and GCaMP6m-X_N_ on DIV 32. (**G**) Cytosolic and nuclear Ca^2+^ oscillations by GCaMP6m-X_C_ (blue) or GCaMP6m-X_N_ (red). In the lower panel, Ca^2+^ traces were rescaled for comparison. (**H**) Key oscillatory characteristics of cytosolic Ca^2+^ by GCaMP6m-X_C_ versus nuclear Ca^2+^ by GCaMP6m-X_N_. (**I** and **J**) The representative two-photon image and spontaneous nuclear Ca^2+^ activities of neurons virally-expressing GCaMP6m-X_N_ in S1 primary somatosensory cortex 5 weeks post virus injection (**I**). The key indices of nuclear Ca^2+^ signals by GCaMP6m-X_N_ (blue), in comparison with GCaMP6m-X_C_ (red) and GCaMP6m (nucleus-filled, cyan) (**J**). (**K** and **L**) Nuclear Ca^2+^ dynamics in S1 primary somatosensory cortex in response to whisker stimulation by *in vivo* two-photon GCaMP6m-X_N_/Ca^2+^ imaging 4 weeks post injection (**K**). Statistical summary of amplitude, FWHM, on rate, off rate and success rate of the trials for GCaMP-X_N_/Ca^2+^ imaging, compared with GCaMP6m-X_C_ (blue) or with nucleus filled GCaMP6m (cyan) (**L**). Standard error of the mean (S.E.M) was calculated when applicable.

Nuclear Ca^2+^ is considered as the fingerprint of electrical activities in neurons (Bading, 2013; Bengtson et al., 2010). Here, GCaMP-X_N_ and GCaMP-X_C_ provided the opportunities to examine the spatiotemporal relationship of cytosolic versus nuclear Ca^2+^ at different stages of neuronal development. By employing GCaMP-X_N_ and GCaMP-X_C_ viruses, we observed that nuclear Ca^2+^ on DIV 32 was able to quickly follow the upstroke of cytosolic Ca^2+^ with prolonged decay phase (***Figure 8F-H*** and ***Figure 8—video supplement 1***). The rather brief lag in time-to-peak between cytosolic Ca^2+^ and nuclear Ca^2+^ indicated that the pathways or mechanisms to elevate nuclear Ca^2+^ are rapid and effective. In comparison, the recovery of nuclear Ca^2+^ (back to the basal level) appears to be a relatively slow process, presumably due to lack of effective Ca^2+^ uptake machinery as compared with cytosolic Ca^2+^. Thus, nuclear Ca^2+^ may fail to exactly follow up cytosolic Ca^2+^ when the intervals between two spikes are too short. In this context, the nucleus tends to function as a type of activity/Ca^2+^ integrator.

Next, we applied GCaMP-X_N_ to mouse brain to examine the nuclear calcium dynamics by *in vivo* two-photon Ca^2+^ imaging. Viral expression of GCaMP6m-X_N_ in S1 primary somatosensory cortex reduced the fluorescence crosstalk from adjacent neuropils, thus providing the desired image contrast sufficient to judge individual neurons by condensed nuclear fluorescence of GCaMP-X_N_ (***Figure 8I***). The nuclear Ca^2+^ dynamics by GCaMP6m-X_N_ showed that the frequency of spontaneous nuclear Ca^2+^ activities *in vivo* was similar to that of cytosolic Ca^2+^ and the FWHM for nuclear Ca^2+^ was much wider than cytosolic Ca^2+^ (***Figure 8J***), resembling the difference between GCaMP-X_C_ and GCaMP-X_N_ *in vitro*. In contrast, the performance of nucleus filled GCaMP6m was much worse than GCaMP-X_N_, as manifested in Ca^2+^ activities of much lower frequency and amplitude. Demonstrated by whisker deflection-evoked nuclear Ca^2+^, the overall performance of GCaMP-X_N_ resembled that of GCaMP6m-X_C_ in the major indices of amplitude, on-rate and success-rate (***Figure 8K, L***). In contrast, the amplitude in average was much smaller for neurons expressing nucleus-filled GCaMP6m than GCaMP6m-X_N_. It is noteworthy that the success-rate of nuclear Ca^2+^ responses was lower than that of cytosolic Ca^2+^ (***Figure 8L***). One possible explanation could be that not all whisker stimuli are transmitted into the nucleus for further signaling and processing. And if true, what is the nature of such selective activation of nuclear signaling cascade? These questions await future investigations by taking advantage of concurrent monitoring of subcellular Ca^2+^, e.g., with both GCaMP-X_C_ and GCaMP-X_N_.

## Discussion

In this work, we applied neuron-compatible versions of GCaMP, including cytosol-localized GCaMP-X_C_ and nucleus-localized GCaMP-X_N_, to monitor Ca^2+^ dynamics across multiple days/weeks both *in vitro* and *in vivo*. Regardless of the delivery methods of transient transfection, viral infection or transgenic expression, GCaMP has intrinsic side-effects accumulated with time, which can be successfully resolved by GCaMP-X. Chronic Ca^2+^ imaging by GCaMP-X not only helped *in vitro* studies to gain unprecedented insights into the roles of subcellular Ca^2+^ in neuritogenesis, but also furnished *in vivo* imaging of live animals with much broader time-window and more reliable Ca^2+^ measurements.

### Solutions currently available to avoid the side-effects of CaM-based GECI

To better utilize CaM-based GECIs *in vitro and in vivo*, solutions with no or minimum side-effects are in need. *In utero* electroporation and viral infection often result in high expression levels particularly near the injection site, and some lines of transgenic mice using weaker promoters could control probe expression within the low levels to alleviate nuclear accumulation (Akerboom et al., 2012; Dana et al., 2019). However, probes present in the cytosol are still problematic, due to the interference with apoCaM-targets represented by CaV1 channels (Yang et al., 2018). Evidently, cytosolic GCaMP affected neural excitability in transgenic mice expressing GCaMP5G or GCaMP6 (Steinmetz et al., 2017). Therefore, it is imperative to improve the conventional CaM-based GECIs, while still inheriting their excellent sensing capabilities in monitoring Ca^2+^ dynamics. One solution is to substitute the key components of GECI design, e.g., to utilize troponin C (from muscle) as a Ca^2+^-binding motif (Mank et al., 2008). The TN-XXL has been claimed to be suitable for chronic imaging potentially benefitted from its origin of muscle (thus less likely to bind endogenous proteins in neurons). However, the TN-XXL solution has at least two shortcomings. First, TNXXL is FRET-based ratiometric sensor, of which the dynamic range is limited by FRET methods and indeed much narrower than GCaMP. Second, the Ca^2+^ binding motif from mammalian troponin C has the canonical EF-hands (resembling CaM), thus still possible to perturb neurons by unexpected binding targets of apo-state, which needs further investigations.

Another approach adopted by this work is to introduce an additional protective motif derived from the IQ domain (CaM-binding domain) of neuromodulin (GAP-43), which is a specific apo-CaM binding motif (Yang et al., 2018). Such apo-CaM binding motif (CBM) is fused onto the N-terminus of conventional GCaMPs (from GCaMP3 to GCaMP6) to obtain new variants of CaM-based GECIs named GCaMP-X. When Ca^2+^ level is low, CBM efficiently prevents apoCaM contained within GCaMP-X from interfering with CaV1 channels and other apoCaM targets (***Figure 1***). Once Ca^2+^ concentration rises, M13 binds to Ca^2+^/CaM with higher affinity than CBM, without jeopardizing Ca^2+^-sensing performance of GCaMP-X inherited from years of efforts. Notably, the perturbation of Ca^2+^/CaM should not be a concern in that the sensor design assures an intramolecular Ca^2+^/CaM binding of high affinity, therefore any perturbation (through intermolecular interactions with endogenous proteins) from Ca^2+^ rise or Ca^2+^/CaM is highly unlikely. In other words, CaM-based GECIs have the intrinsic protection for Ca^2+^ conditions but not for the apo state, the latter of which should be focused on to avoid the side-effects of future GECIs. In fact, GCaMP-X is no longer accumulated in the nucleus even without any localization tag, further supporting that apoCaM protection (from binding to the channel) is critical to cytosol-nuclear translocation thus alleviating nuclear filling (Yang et al., 2018). Collectively, these considerations or rules centered with apoCaM/Ca^2+^-CaM binding should be broadly applicable to CaM-based sensors or actuators (Grødem et al., 2021; Haiech et al., 2019).

### Spontaneous Ca^2+^ activities in association with neuronal development

While the membrane voltage is oscillating, cellular and subcellular Ca^2+^ signals are also fluctuating, closely involved in neuronal development and circuit formation both *in intro* and *in vivo* (Kirkby et al., 2013; Sun et al., 2010). Meanwhile, in line with the “calcium hypothesis”, dysregulated Ca^2+^ oscillations would lead to defective morphology and functions of neurons, and eventually neural diseases (Khan et al., 2020). In Ca^2+^ imaging experiments, Ca^2+^ fluorescence signals and electrical activities are often referred to each other due to the fact that action potentials initiated by Na^+^ channels would subsequently drive the fast opening of Ca^2+^ channels to induce Ca^2+^ influx. Thus, MEA (multiple electrode array) as an electrical recoding platform is widely used in long-term live-cell monitoring *in vitro* (Obien and Frey, 2019; Shafer, 2019). On the other hand, membrane potentials induced by Na^+^ or Ca^2+^ influx could have distinguished mechanisms and consequences, which is also true for Ca^2+^ oscillations of different forms, e.g., subthreshold oscillations by L-type Ca^2+^ channels, or intracellular Ca^2+^ fluctuations by intracellular Ca^2+^-release channels (Chan et al., 2007; Uhlen and Fritz, 2010). GECIs are the ideal alternative tools to overcome the limitations of electrical methods such as MEA, if the cell compatibility issues are resolved as demonstrated by GCaMP-X. GECI imaging methods directly and faithfully capture Ca^2+^ activities at different loci in the cell, allowing high spatial-temporal resolution of concurrent morphological/functional imaging. Subcellular Ca^2+^ oscillations may be responsible for different aspects of neurogenesis and neuritogenesis. In fact, MEA-based data suggest that spontaneous Ca^2+^ activities and neuronal development are correlated with each other (Obien and Frey, 2019; Van et al., 2004; Yada et al., 2016), consistent with Ca^2+^-dependent mechanisms of morphogenesis (Kater et al., 1994; Kater et al., 1988). Chronic imaging experiments with GCaMP-X in this work have provided the platform for further investigations into the temporal/spatial details on Ca^2+^-dependent morphogenesis. Earlier Ca^2+^ activities (of higher frequency) may represent spontaneous activity before synapse or network formation (Spitzer, 2006). At the later stage, synchronized Ca^2+^ oscillations (of lower frequency) emerge, along with dramatic changes in neural development. Autonomous oscillations of SNc neurons switch from HCN channel-based to CaV1.3 channel-based in Parkinson’s disease or neural aging (Chan et al., 2007). Cultured cortical slices and hiPSC-derived cortical neurons also suggest that L-type Ca^2+^ channels are crucial for both spontaneous Ca^2+^ activities and neuronal development (Horigane et al., 2020; Plumbly et al., 2019). A similar mechanism is likely shared by the spontaneous/oscillatory Ca^2+^ activities observed in this study (***Figure 3*** and ***Figure 4***), which awaits in-depth investigations in the future. Moreover, the expression, trafficking and functions of ion channels and receptors are also subject to regulations by activities of different patterns (Ruffinatti et al., 2013; Spitzer, 2006; Toth et al., 2016), for which chronic GECI imaging should be of great help to elucidate these compound effects and mechanisms. In this work, we have particularly focused on spontaneous Ca^2+^ activities of cortical neurons in association with neuronal development both *in vitro* (***Figure 3*** and ***Figure 4***) and *in vivo* (***Figure 2***, ***Figure 5*** and ***Figure 6***), which serves as the exemplars of chronic GCaMP-X imaging. Importantly, if such Ca^2+^-neuritogenesis coupling is perturbed by GCaMP, the side-effects would be induced in neurons as supported by data from both *in vitro* and *in vivo*.

### Improved neuron-compatibility of GCaMP-X_C_ and -X_N_

GCaMP, as widely-applied GECI, has been evolved into its 8^th^ version (Grødem et al., 2021), with enhanced sensitivity, brightness and kinetic properties tailored to specific imaging purposes, including single action potentials and activities in neuronal populations and microcompartments (Dana et al., 2019). However, the cytotoxicity of GCaMP has been a persistent problem from early on, mainly due to the fact that the CaM-centered schemes of GCaMP have been largely inherited across generations of its design (Akerboom et al., 2012; Chen et al., 2013; Dana et al., 2019; Nakai et al., 2001; Tallini et al., 2006; Tian et al., 2009). *In vivo* Ca^2+^ imaging with GCaMP viruses is facing the dilemma of safety versus reliability. On one hand, reducing GCaMP levels could alleviate or postpone the cytotoxicity, but low levels of expression would also reduce the contrast and SNR (Rose et al., 2014). On the other hand, increasing the expression level of GCaMP would enhance the data quality of Ca^2+^ imaging, but exacerbate the damages to neurons (Resendez et al., 2016). GCaMP transgenic mice are encountered with similar problems. Several lines of GCaMP transgenic mice reported earlier, such as Emx1-Cre;Ai38 GCaMP3 transgenic mice, attempted to resolve the safety issue by restricting the expression to ultra-low level (∼5 μM) (Zariwala et al., 2012), but sacrificing the imaging quality (Rose et al., 2014). The mouse lines reported recently, such as Emx1-Cre;CaMK2*α*-tTA;Ai93 GCaMP6f transgenic mice and Slc17a7-IRES2-Cre;Ai148 GCaMP6f transgenic mice (Daigle et al., 2018; Madisen et al., 2015), managed to elevate the expression levels. However, epileptiform activities have been observed from these mice presumably due to perturbations of Ca^2+^ signaling observed *in vitro* (Daigle et al., 2018; Steinmetz et al., 2017). One bypass solution is to conditionally induce GCaMP expression in neurons to conduct GCaMP imaging within a time window (shortly after induction of GCaMP expression), but long-term expression and/or chronic imaging still suffer from the problematic side-effects. Instead, GCaMP-X allows long-term and high-level expression to increase the quality and reliability of Ca^2+^ measurement while preventing the aforementioned side-effects on neurons (Yang et al., 2018). For *in vitro* studies, investigations of long-term Ca^2+^ dynamics are largely hindered by the cytotoxicity of GECIs or dyes (Rose et al., 2014; Smith et al., 2018). GCaMP-X is well suited to study Ca^2+^ dynamics during neural development, or for high-throughput fluorescence assays to screen compounds with long-term effects (Vetter et al., 2020). For *in vivo* studies, due to the aforementioned concerns in GCaMP applications (Resendez et al., 2016), false-negative or false-positive results by nuclear-filled GCaMP or under-expressed GCaMP are misleading especially in imaging large populations of neurons. GCaMP-X provides *in vivo* experiments with higher SNR, better precision, lower neurotoxicity and longer term of expression/imaging.

CaM-based GECIs with nuclear-localization tags cause serious damage in both neuronal development and Ca^2+^ signals *in vitro* and *in vivo* (Chen et al., 2020; Shemesh et al., 2020; Yang et al., 2018). In contrast, neuron-compatible GCaMP-X_N_ allows to directly monitor nuclear Ca^2+^ dynamics, an important aspect of intracellular calcium signaling (Bading, 2013). By combining GECIs specific to different compartments including cytosolic GCaMP-X_C_ and nuclear GCaMP-X_N_, a complete set of new information could be acquired (***Figure 8***), e.g., nuclear Ca^2+^ signals in relation to neuronal activities (Bading, 2013; Bengtson et al., 2010). Considering that nuclear Ca^2+^ signals directly regulate transcription and gene expression of neurons (Hardingham et al., 2001; Hardingham et al., 1997), Ca^2+^ dynamics in the nucleus might play more sophisticated roles beyond simple fingerprints of cytosolic Ca^2+^. There have been discussions about the generation of nuclear Ca^2+^ signals independent of cytosolic Ca^2+^ (Echevarria et al., 2003; Marchenko et al., 2005). As nucleus-compatible GECI, GCaMP-X_N_ teamed up with other subcellular (multi-color) probes should help address these intriguing questions (Inoue et al., 2015; Inoue et al., 2019). In addition, soma-targeted ribo-GCaMP and somaGCaMP have been recently reported, which restrict the expression of GCaMP to the soma in order to reduce neuropil crosstalk and increase the signal-to-noise ratio (Chen et al., 2020; Shemesh et al., 2020). An alternative solution to reduce the crosstalk is to restrict GCaMP in the nucleus (Kim et al., 2014; Nguyen et al., 2016), which should enhance the spatial precision of imaging/analyzing individual neurons. Unfortunately, NLS-targeted GCaMP exhibits slow and small responses to stimuli (Chen et al., 2020; Shemesh et al., 2020), due to its perturbations on apoCaM-binding proteins (Jurado et al., 1999; Yang et al., 2018). In this regard, GCaMP-X_N_ should have similar or even better performance in comparison with ribo-GCaMP and somaGCaMP, which awaits future investigations.

### Design and application guidelines for CaM-based GECIs

Base on the previous works and this study, we here summarize the guidelines to choose proper probes and/or experimental conditions for chronic Ca^2+^ imaging applications with CaM-based GECIs. First, if the candidate probes have known (chronic) side-effects, imaging should be performed as soon within the safety window. Second, it is preferable to pick those probes with high levels of brightness, dynamic-range and SNR, since it would reduce expression time/level and thus help alleviate any cellular damage. Third, if the probes with intrinsic toxicity have to be selected, additional precautions are expected such as conditional/controlled expression or weak promoters, etc. For the design of GECIs based on CaM or similar sensing elements, several rules or concerns have been learned thus far: 1) avoid any free motif or key element to stay unbound at their apo or basal states; 2) employ high-affinity interactions within the intermolecular pair; 3) choose the template genes of remote origins in terms of cell types, tissues, and evolutionary relationships, to avoid potential interference in the first place; 4) minimize the functional motif(s) deployed by the design and include the protection strategy when necessary; 5) carefully test the probes *in vitro* at the early phase of design procedures, by mimicking physiological or *in vivo* conditions. These strategies in choosing and devising CaM-based GECIs could also be extended onto other molecular tools based on CaM binding or even protein-protein interactions in general.

## Methods

### Molecular biology

pGP-CMV-GCaMP6m (#40754), pGP-CMV-jGCaMP7b (#104484), pEGFP-N1-GCaMP6m-X_C_ (#111543), GCaMP6m-X_N_ (#118976) and AAV-*Syn*-GCaMP6m-X_C_ (#118975) are available on Addgene. GCaMP7b-X_C_ and GCaMP7b-X_N_ were constructed by replacing previously reported GCaMP6m-X_C_ or GCaMP6m-X_N_ (Yang et al., 2018) with appropriate PCR-amplified segments from jGCaMP7b via unique EcoRI / HindIII sites, or EcoRI / NotI sites, respectively. pEGFP-N1-GCaMP7b-X_C_ (#178361) and pEGFP-N1-GCaMP7b-X_N_ (#178362) have been deposited onto Addgene.

### Mice

Procedures involving animals have been approved by local institutional ethical committees (IACUC in Tsinghua University and Beihang University). *In vivo* experiments were based on data from 21 mice (age>P60, both male and female). Thirteen C57BL/6 mice were used for expression tests and functional tests of three Ca^2+^ indicators (GCaMP6m, GCaMP6m-X_C_ and GCaMP6m-X_N_) and two C57BL/6 mice were used for GFP virus injection. Six Ai148D x Rasgrf2-2A-dCre (Jax #030328; Jax #022864) mice were used for the tests of GCaMP6f-transgenic mice. Expression of GCaMP6f in Rasgrf2-2A-dCre;Ai148D mice was induced with antibiotic Trimethoprim (TMP) by intraperitoneal injection with the dose of 0.25-0.5 mg/g *in vivo* (Sando et al., 2013). *In vitro* experiments were based on data from 53 mice (P0-P1, both male and female). 39 ICR mice were used for expression and functional tests of Ca^2+^ indicators. Two newborn Ai148D x Emx1-Cre mice were used to persistently express GCaMP in cultured neurons. In the tests of cortex neurons expressing GCaMP6f from Rasgrf2-2A-dCre;Ai148D mice, nine positive Rasgrf2-2A-dCre;Ai148D mice were compared with three negative-control mice, by directly adding 10 μM TMP (to induce GCaMP6f expression) into growth medium after dissection.

### Dissection and culturing of cortical neurons

Cortical neurons were dissected from newborn ICR mice or Ai148 GCaMP6f-transgenic mice. Isolated tissues of cortex were digested with 0.25% trypsin for 15 min at 37 °C. Then digestion was terminated by DMEM supplemented with 10% FBS. The cell suspension was sieved through a filter and centrifuged at 1000 rpm for 5 min. The cell pellet was resuspended in DMEM supplemented with 10% FBS and were plated on poly-D-lysine-coated 35 mm No. 0 confocal dishes (In Vitro Scientific). After 4-6 hours, neurons were maintained in Neurobasal medium supplemented with 2% B27 and 1% glutaMAX-I (growth medium), and cultured in the incubator with temperature of 37 °C and 5% CO_2_. Fresh growth medium was supplemented to neurons every 3-4 days to maintain the volume of 2 ml growth medium. All animals were obtained from the laboratory animal research center, Tsinghua University. Procedures involving animals has been approved by local institutional ethical committees (IACUC in Tsinghua University and Beihang University).

### Virus infection on cultured neurons

Viruses of GCaMP6m (AAV2/DJ-*Syn*-GCaMP6m), GCaMP6m-X_C_ (AAV2/DJ-*Syn*-GCaMP6m-X_C_), GCaMP6m-X_N_ (AAV2/DJ-*Syn*-GCaMP6m-X_N_), jGCaMP7b (AAV2/DJ-Syn-jGCaMP7b) and jGCaMP7b-X_C_ (AAV2/DJ-Syn-jGCaMP7b-X_C_) were provided by Hanbio Biotechnology, China. The neuron broad-spectrum promoter Syn and AAV2/DJ serotypes were selected for neuro-specific expression of GCaMP or GCaMP-X in cultured cortical neurons. 1-2 μl 1x10^12^ v.g./ml of the three kinds of viruses were added to growth medium on DIV 0, respectively. The expression of GCaMP and GCaMP-X was detectable on DIV 3, reached the peak on DIV 7 and sustained the high-level up to one-month.

### Ca^2+^ imaging with GCaMP or GCaMP-X in cultured cortical neurons

Ca^2+^ imaging of neurons expressing GCaMP or GCaMP-X was acquired by confocal microscopy (A1RMP, Nikon, Japan; Dragonfly 200, Andor, England). 488 nm laser was used for excitation. 35 mm confocal dish containing cultured cortical neurons was set in the live-cell imaging culture chamber of the confocal microscope to maintain the environment of 37°C, 5% CO_2_ and 95% humidity. Sampling rate of images was at 1-5 Hz and 3-5 view fields were selected from each dish. Fluorescence intensity (*F*) was subtracted from its background. *F0* is the baseline fluorescence averaged from 5 data points at rest, and Δ*F* = *F*-*F0*. Δ*F/F0* serves as the index for Ca^2+^ dynamics. Ca^2+^ waveforms were analyzed by the gadget of Quick Peaks in Origin software with the Three-Standard-Deviations Rule (values > 3 S.D.). On and off rate were characterized by the time to rise up or decay down to 50% of the maximum (Δ*F/F0*), respectively. And the full width at half maximum (FWHM) is defined as the duration of time between the (upward and downward) half-maximum timepoints. Mean of correlation coefficients was evaluated by Spearman Rank Correlation Coefficient in Origin software. FFT analyses were performed in Origin software. One-sided is selected for spectrum type and amplitude is define as below:

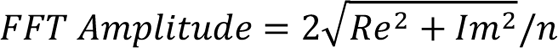

Here *Re* and *Im* are the real and imaginary parts of FFT results, and *n* is the size of the input signal.

Neurite tracings were depicted with Imaris 7.7.2 from the images under average intensity projection. Morphological analysis and Ca^2+^ oscillation analysis were performed with at least 30 neurons at each time point in each independent experiment. Fluorescence intensities of Ca^2+^ dynamics in neurons were color-coded by Matlab (Mathworks) and Fiji.

### Transfection, confocal fluorescence imaging and analysis of neurite morphology

2 μg of cDNA encoding jGCaMP7b, jGCaMP7b-X_C_ or jGCaMP7b-X_N_ and 1 μg cDNA encoding CFP (for labelling the soma area and neurites) were transiently transfected into DIV 5-7 cultured cortical neurons by Lipofectamine 2000 (Invitrogen) with a typical protocol according to the manual. The opti-MEM containing plasmids and Lipofectamine 2000 was added to the Neurobasal medium for transfection. After 2 hours, neurons were maintained in Neurobasal medium supplemented with 2% B27, 1% glutaMAX-I for at least 2 days before analyzing neurite morphology.

Fluorescence imaging of cultured cortical neurons was performed on ZEISS Laser Scanning Confocal Microscope (LSM710, Carl Zeiss) and ZEN 2009 software. N/C ratio of GCaMP or GCaMP-X was calculated by the ratio of fluorescence intensity (nuclear/cytosolic). Measurement of the total length and *Sholl* analysis for neurites were performed with Imaris 7.7.2 (Bitplane).

Only nonoverlapping neurons were selected for analysis and images of at least 24 neurons from two independent culture preparations were analyzed. Neurite tracings were depicted with Imaris 7.7.2 in CFP channel.

### Craniotomy and *in vivo* virus injection

Wildtype mice were used for virus injection and craniotomy under isoflurane anesthesia (5% for induction, 1-1.5% during surgery). AAV2/9-*Syn*-GCaMP6m-X_C_ and AAV2/9-*Syn*-GCaMP6m-X_N_ (1.0x10^13^ v.g./ml, customized by BrainVTA, Wuhan, China) viruses were tested in the primary somatosensory cortex (S1BF: AP -1.5, ML -3.0, DV 0.2/0.4) with AAV2/9-*Syn*-GCaMP6m (1.0x10^12^ v.g./ml, BrainVTA, Wuhan, China) virus as a control. AAV2/1-CAG-GFP (4.7x10^12^ v.g./ml, UNC) was injected as a control for soma size analysis in ***Figure 7A***.

Craniotomy was done three weeks after virus injection. A piece of skull above S1BF was removed to expose a square imaging window (∼3 mm x 3 mm, centered on S1BF) and the cortex was protected by a hand-cut glass window using #1 coverglass. Then the glass window was fixed using adhesive (Krazy glue, Elmer’s Products Inc) and dental cement. A head-post was also fixed to the posterior area of the mouse head using dental cement. 0.2ml flunixin meglumine (0.25 mg/ml) was subcutaneously injected after surgery for 3 consecutive days.

### Preparations of brain slices

Mice were anaesthetized by intraperitoneal injection of avertin solution (250-500 mg/kg). Then they were transcardially perfused with phosphate buffer saline (PBS) followed by 4% Paraformaldehyde (PFA) solution. Brains were immersed in 4% PFA solution overnight and were embedded in 2.5% agarose gel for slicing operation. Slices were obtained using the Lecia vibratome (LeciaVT1200S) with proper parameters including depth, speed and thickness of the brain section (70 or 100 µm).

### *In vivo* two-photon Ca^2+^ imaging to monitor spontaneous Ca^2+^ oscillation and Ca^2+^ dynamics under whisker test

A 2-photon random access mesoscope controlled with ScanImage 2017 (Vidrio Technologies) was used for *in vivo* Ca^2+^ imaging (Pologruto et al., 2003; Sofroniew et al., 2016). Images (512x512 pixels, 600x600 or 300x300 μm^2^) of L2/3 cells (150-250 μm under pia) in S1BF were collected at 7.4 Hz frame rate. Laser power (970 nm) was up to 60 mW out of objective. Calcium signal was extracted using CaImAn toolbox and data analysis was performed using Matlab (Giovannucci et al., 2019).

For functional test, imaging was carried out together with contralateral whisker stimulation using a 1.2 mm-diameter pole (3.16 mm in amplitude, 10 vibrations in 0.5 s or 1 s for each trial). For each ROI, 20-40 trials were performed and calcium signaling was aligned to the onset of the whisker stimulation.

The fluorescence of each neuron was measured by averaging all pixels within the ROI (regions of interest) and corrected for neuropil contamination. The fluorescence signal was estimated by the equation below:

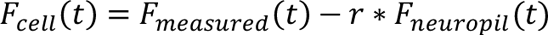

where *r*=0.7 and *F_neuropil_(t)* was measured by averaging the fluorescence signal of all pixels within a 40 μm radium from the cell (Chen et al., 2013).

### Whole-cell electrophysiology

HEK293 cells (ATCC) were cultured in 60 mm dishes and checked by PCR with primers 5’-GGCGAATGGGTGAGTAACACG -3’ and 5’-CGGATAACGCTTGCGACCTATG -3’ to ensure free of mycoplasma contamination. Recombinant channels (α_1DL_, β_2a_ and α_2δ_ subunits, 4 μg each) were transiently transfected according to established calcium phosphate protocol (Liu et al., 2017a; Liu et al., 2017b; Liu et al., 2010). To enhance expression, cDNA for simian virus 40 T antigen (1 µg) was also co-transfected. Additional 2 µg of cDNA of GCaMP7b or GCaMP7b-X_C_ was added as required in co-transfections. Whole-cell recordings of transfected HEK293 cells were performed at room temperature (25 °C) using an Axopatch 200B amplifier (Axon Instruments). The internal solutions contained, (in mM): CsMeSO_3_, 135; CsCl_2_, 5; MgCl_2_, 1; MgATP, 4; HEPES, 5; and EGTA, 5; at 290 mOsm adjusted with glucose and at pH 7.3 adjusted with CsOH. The extracellular solution contained (in mM): TEA-MeSO_3_, 135; HEPES, 10; CaCl_2_ or BaCl_2_, 10; 300 mOsm, adjusted with glucose and at pH 7.3 adjusted with TEAOH. Whole-cell currents were generated from a family of step depolarizations (-70 to +50 mV from a holding potential of -70 mV and step of 10 mV).

### Data analysis and statistics

Data were analyzed in Matlab, OriginPro and GraphPad software. Standard error of the mean (S.E.M.) and two-tailed Student’s *t*-test or one-way ANOVA followed by Bonferroni for post hoc tests were calculated when applicable. Criteria of significance: *, *p* < 0.05; **, *p* < 0.01; ***, *p* < 0.001; and *n.s.* denotes “not significant”. All experiments were performed at least twice with appropriate sample sizes.

## Acknowledgements

We thank all X-Lab members for discussions and help. This work is supported by grants from Natural Science Foundation of China (NSFC 21778034, 81971728 and 11902021) and of Beijing (BNSF 7191006 and 5204037), China Postdoctoral Science Foundation (BX20180027 and 2018M641146), and open fund from Laboratory for Biomedical Engineering of Ministry of Education, Zhejiang University.

## Author Contributions

XDL, ZVG and YXY conceived and supervised the project; JLG, WXL, YJT, YMG, YTL and YXY performed the experiments; JLG, WXL, YXY, YMG, YJT and XDL analyzed the data; XDL, WXL, GJL and YXY wrote the manuscript; all authors participated in reviewing and commenting on the manuscript.

---

**Figure 1—supplement 1.**
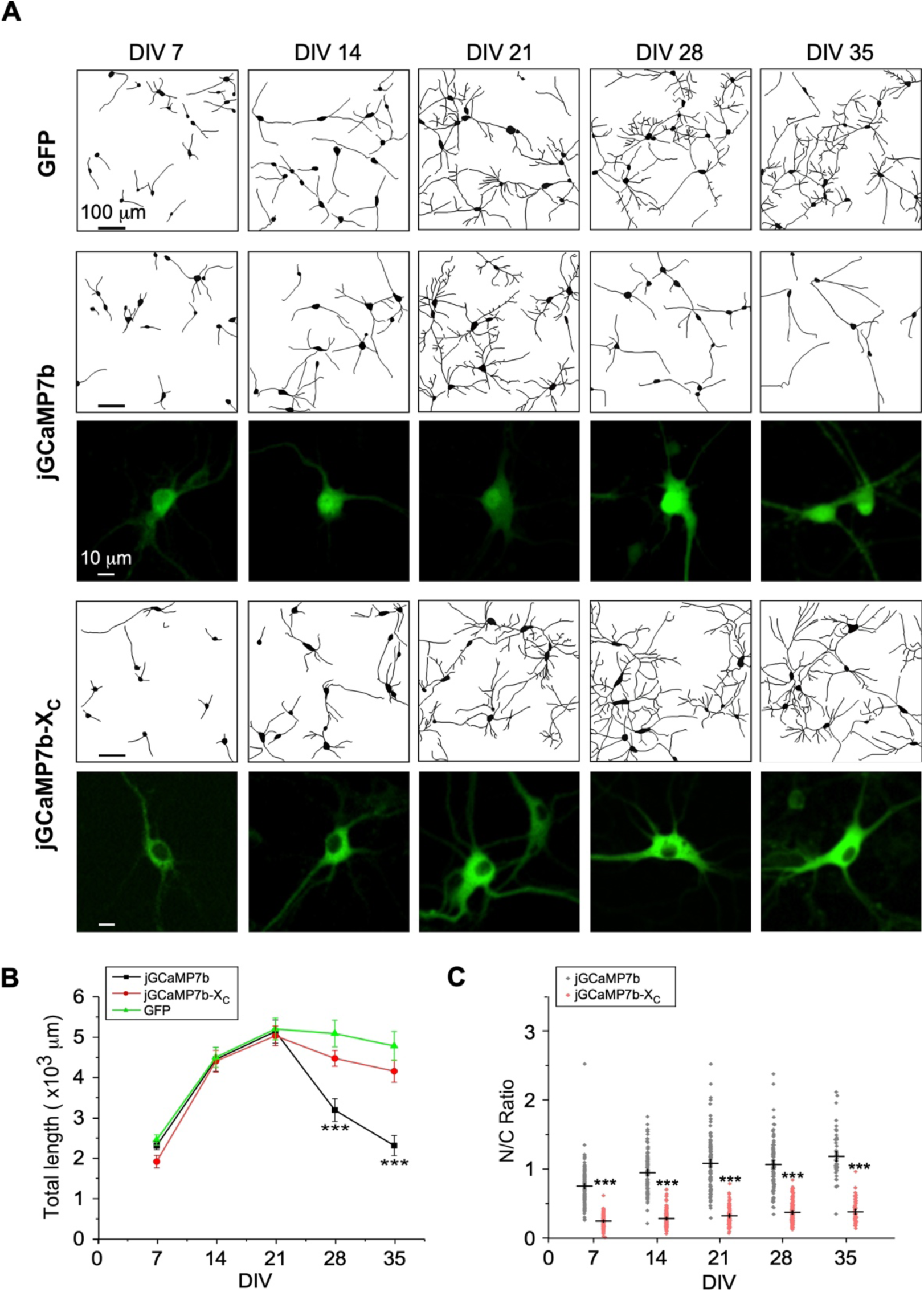
Morphological analysis for cultured cortical neurons. (**A**) Cortical neurons virally-expressing jGCaMP7b-X_C_, jGCaMP7b and GFP control, respectively. For each group, neurite tracing of multiple neurons (upper rows) and representative confocal fluorescence images of individual neurons (lower rows) at the selected timepoints of DIV 7 to DIV 35. (**B**) Temporal profiles of the total length of neurites summarized for jGCaMP7b-X_C_ and jGCaMP7b versus GFP control. (**C**) N/C ratio of neurons expressing jGCaMP7b or jGCaMP7b-X_C_. Standard error of the mean (S.E.M) and Student’s *t*-test (two-tailed unpaired with criteria of significance: *** *p*<0.001) were calculated when applicable.

**Figure 3—figure supplement 1.**
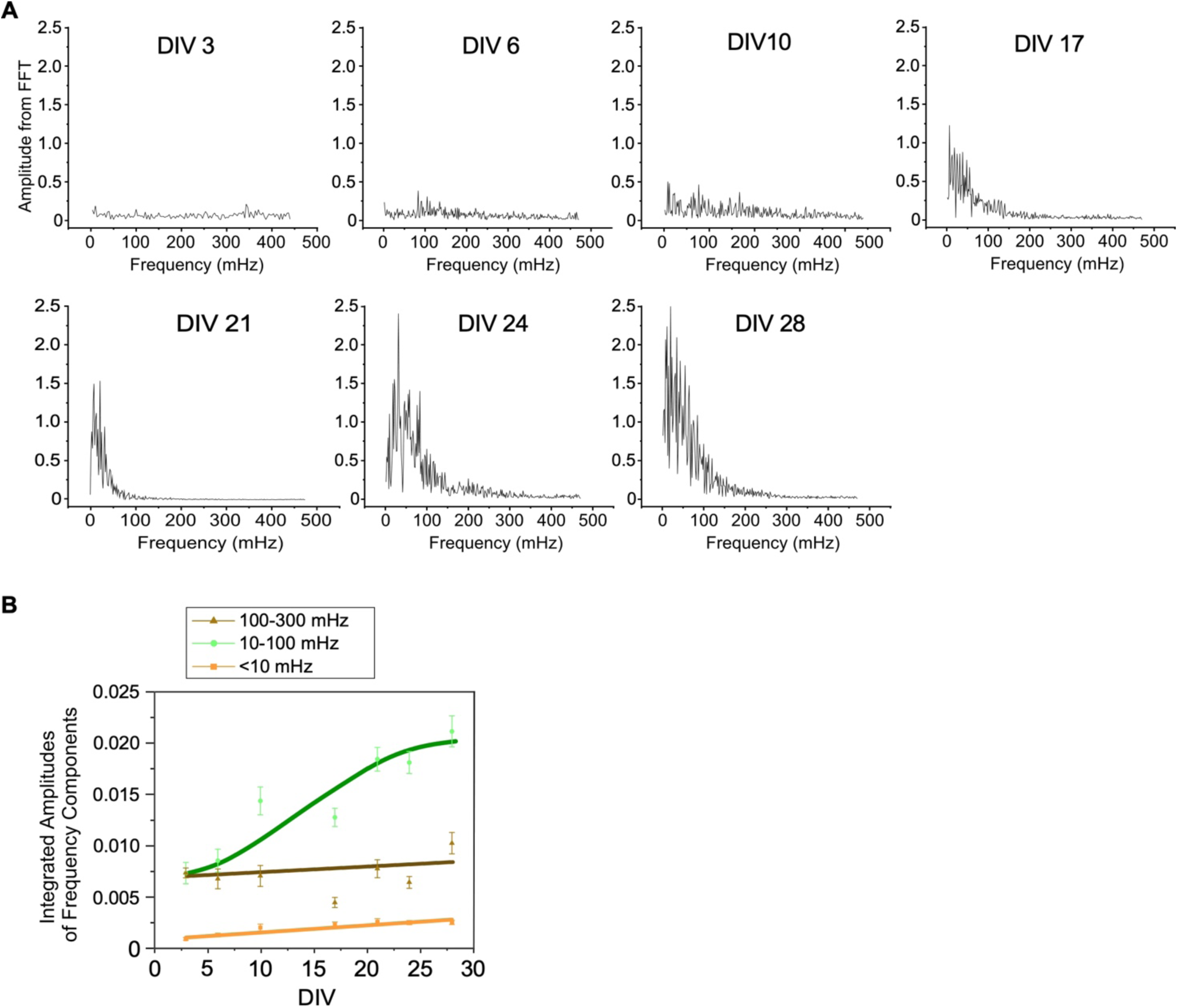
Spectral analysis for Ca^2+^ waveforms acquired by GCaMP-X. (A) Representative FFT-based power spectra of individual neurons at different timepoints (DIV). Neurons from DIV 3 to DIV 10 exhibited low FFT amplitudes, whereas neurons from DIV 21 to DIV 28 showed significantly large amplitudes with the central frequency around 10-100 mHz. DIV 17 appeared to be the transition time for neurons to develop from weak oscillations (DIV 3 to DIV 10) to strong oscillations (DIV 21 to DIV 28). (**B**) Temporal profiles for the major frequency components of Ca^2+^ oscillations. Three frequency components including 100-300 mHz (fast), 10-100 mHz (slow) and <10 mHz (ultraslow) were summarized (n=3 individual experiments). Frequency components were evaluated by the amplitude integration over each frequency band in FFT spectra.

**Figure 3—supplement 2.**
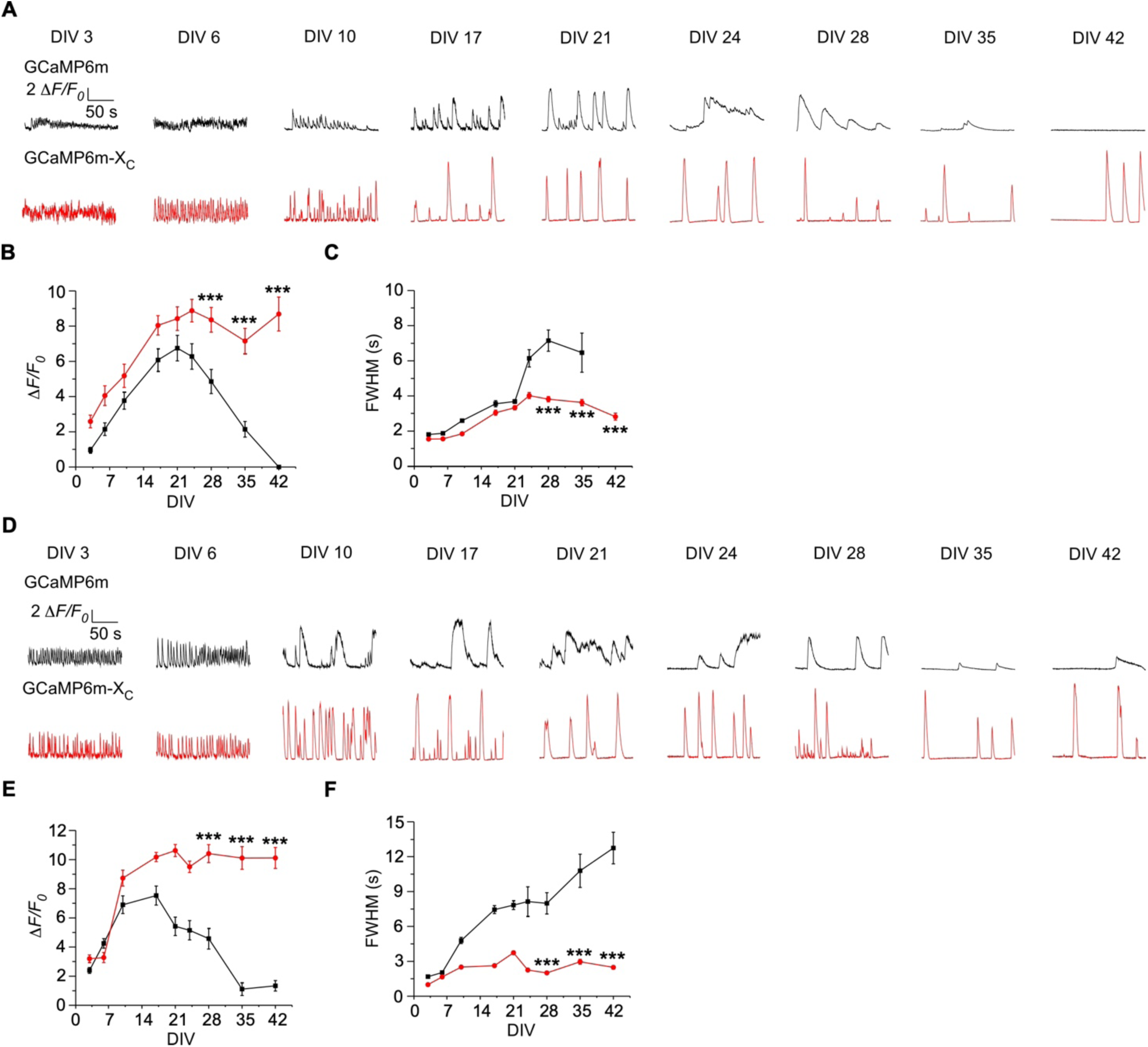
Ca^2+^ oscillations in long-term cultured cortical neurons expressing GCaMP6m or GCaMP6m-X_C_ *in vitro*. (**A-C**) Ca^2+^ activities in cultured cortical neurons expressing GCaMP6m (black) or GCaMP6m-X_C_ (red), respectively. Exemplar traces from DIV 3 to DIV 42 (**A**), summarized by peak amplitudes (Δ*F/F0*, **B**) and full width at half maximum (FWHM, **C**). Note that GCaMP6m on DIV 42 did not record any noticeable response. (**D**-**F**) Another set of Ca^2+^ oscillation data and analyses over multiple weeks similar to (**A**-**C**). Standard error of the mean (S.E.M) and Student’s *t*-test (two-tailed unpaired with criteria of significance: *** *p*<0.001) were calculated when applicable.

**Figure 3—supplement 3.**
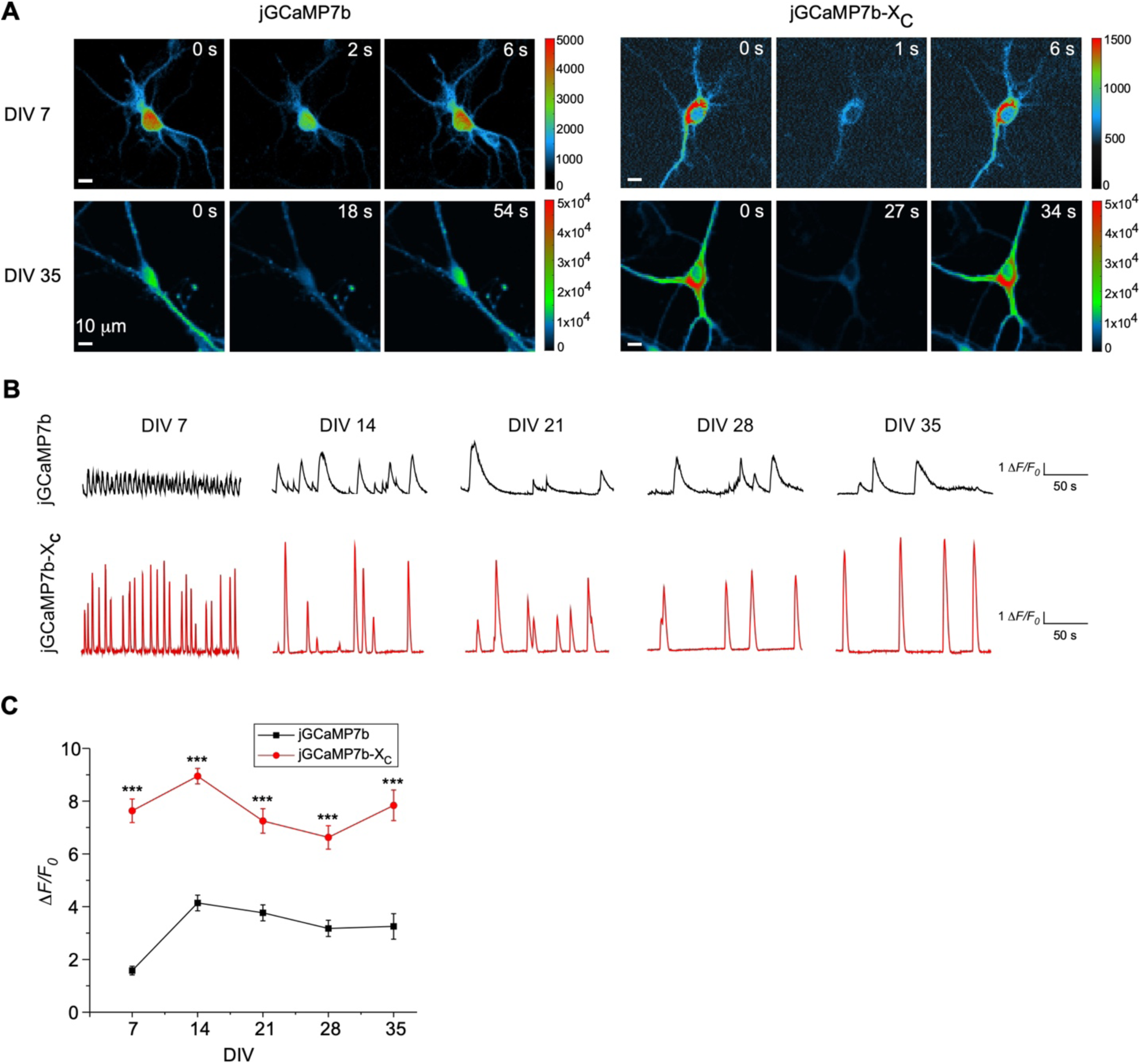
Ca^2+^ oscillations in cultured cortical neurons expressing jGCaMP7b or jGCaMP7b-X_C_. (**A**) Representative Ca^2+^ fluorescence images (color-coded) of cultured cortical neurons expressing jGCaMP7b or jGCaMP7b-X_C_ on DIV 7 or DIV 35. (**B**) Representative Ca^2+^ activity traces of cultured cortical neurons expressing jGCaMP7b (black) or jGCaMP7b-X_C_ (red) across 5 weeks (from DIV 7 to DIV 35). (**C**) Summarized peak amplitudes in time-dependent profiles (*ΔF/F0*). Standard error of the mean (S.E.M) and Student’s *t*-test (two-tailed unpaired with criteria of significance: *** *p*<0.001) were calculated when applicable.

**Figure 4—supplement 1.**
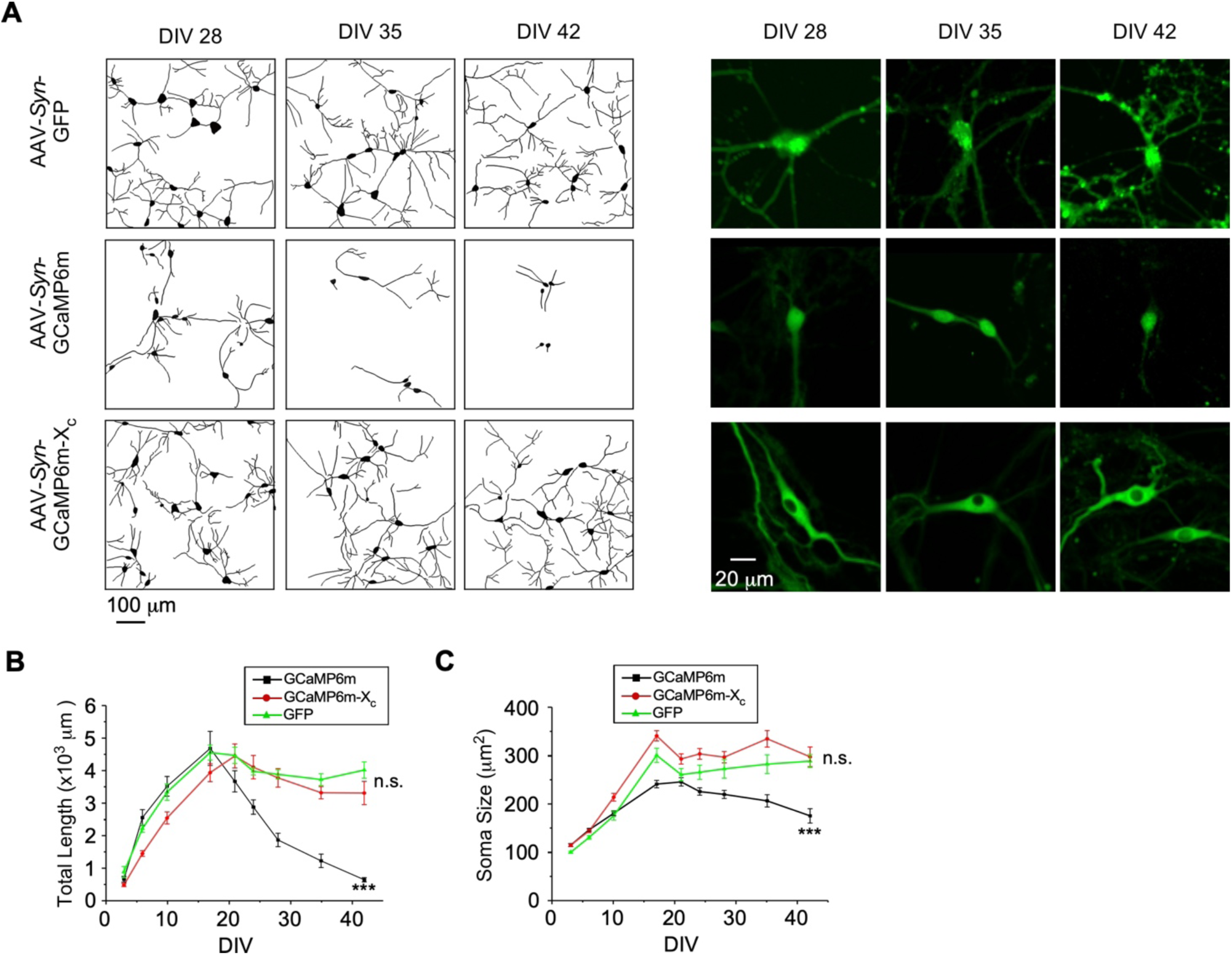
Indistinguishable neuronal development between GCaMP6m-X_C_ and GFP of long-term expression. (**A**) Representative neurite tracing of multiple neurons (left) and fluorescence images of individual neurons (right) for cultured cortical neurons expressing GCaMP6m-X_C_, GCaMP6m or GFP on DIV 28, DIV 35 and DIV 42. (**B** and **C**) Temporal profiles of the total length of neurites (**B**) and the soma size from DIV 3 to DIV 42 (**C**). Compared on DIV 42, no significant difference exists between GCaMP6m-X_C_ and GFP control; and the differences between GCaMP6m-X_C_ and GCaMP6m are significant. Standard error of the mean (S.E.M) and Student’s *t*-test (two-tailed unpaired with criteria of significance: * *p*<0.05, ** *p*<0.05 and *** *p*<0.001) were calculated when applicable, and n.s. denotes “not significant”.

**Figure 4—figure supplement 2.**
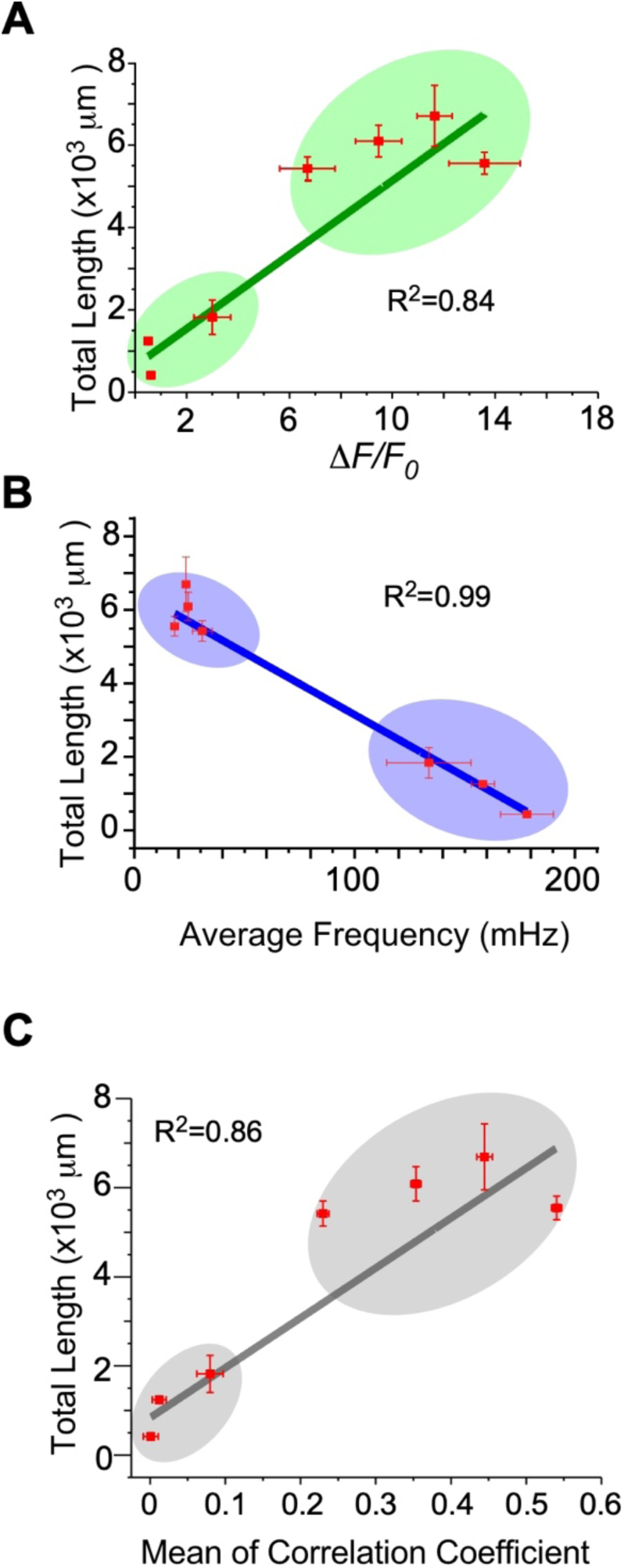
Potential relationships between and neurite length and oscillation characteristics. The peak amplitude (**A**), the average frequency (**B**), or the level of synchrony (**C**) was examined for its potential correlation with the total length of neurites, based on the data acquired by GCaMP6m-X_C_ in Figure 3 and Figure 4. The linear correlation coefficient (*R^2^*) was evaluated between each index and neurite length.

**Figure 7—figure supplement 1.**
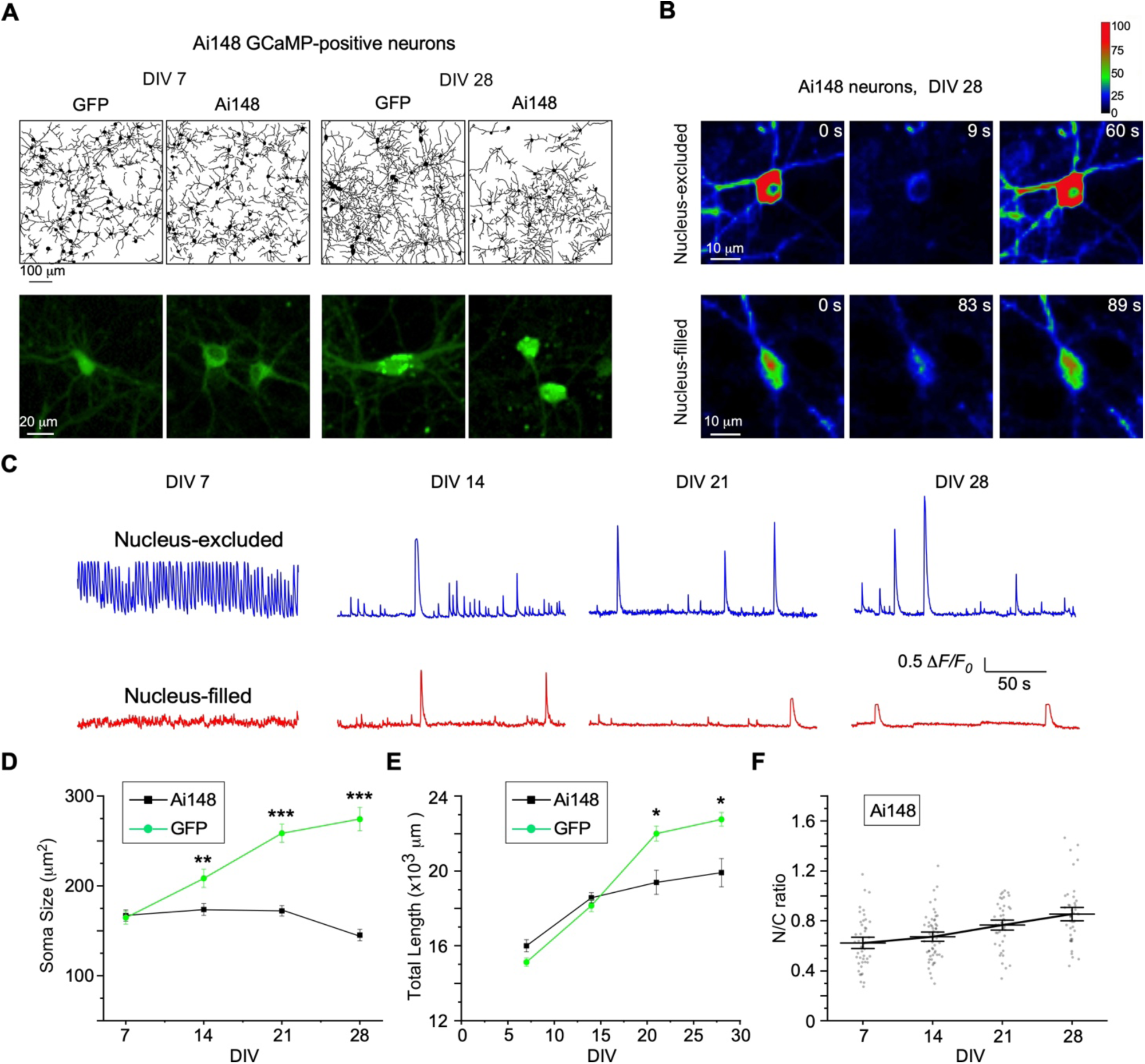
Chronic evaluation of cultured cortical neurons from Ai148 mice with inducible GCaMP6f expression. (**A**) Representative neurite tracing and fluorescence images for cultured cortical neurons expressing GCaMP6f or GFP on DIV 7 and DIV 28, respectively. GCaMP6f expression was induced by 10 μM TMP from Rasgrf2-2A-dCre x Ai148 mice. GFP control neurons were also treated with the same dose of TMP. (**B** and **C**) Spontaneous Ca^2+^ oscillations from nucleus-excluded versus nucleus-filled neurons (with the criteria of N/C ratio<0.6 or >0.8, respectively) from Ai148 mice, shown by sampled images at particular timepoints as indicated (**B**). Representative Ca^2+^ waveforms from long-term GCaMP imaging from DIV 7 to DIV 28 (**C**). (**D-F**) Temporal profiles of soma size (**D**), total neurite length (**E**), and N/C ratio of GCaMP6f (**F**) for Ai148 neurons. Standard error of the mean (S.E.M) and Student’s *t*-test (two-tailed unpaired with criteria of significance: * *p*<0.05, ** *p*<0.01, *** *p*<0.001) were calculated when applicable.

**Figure 7—figure supplement 2.**
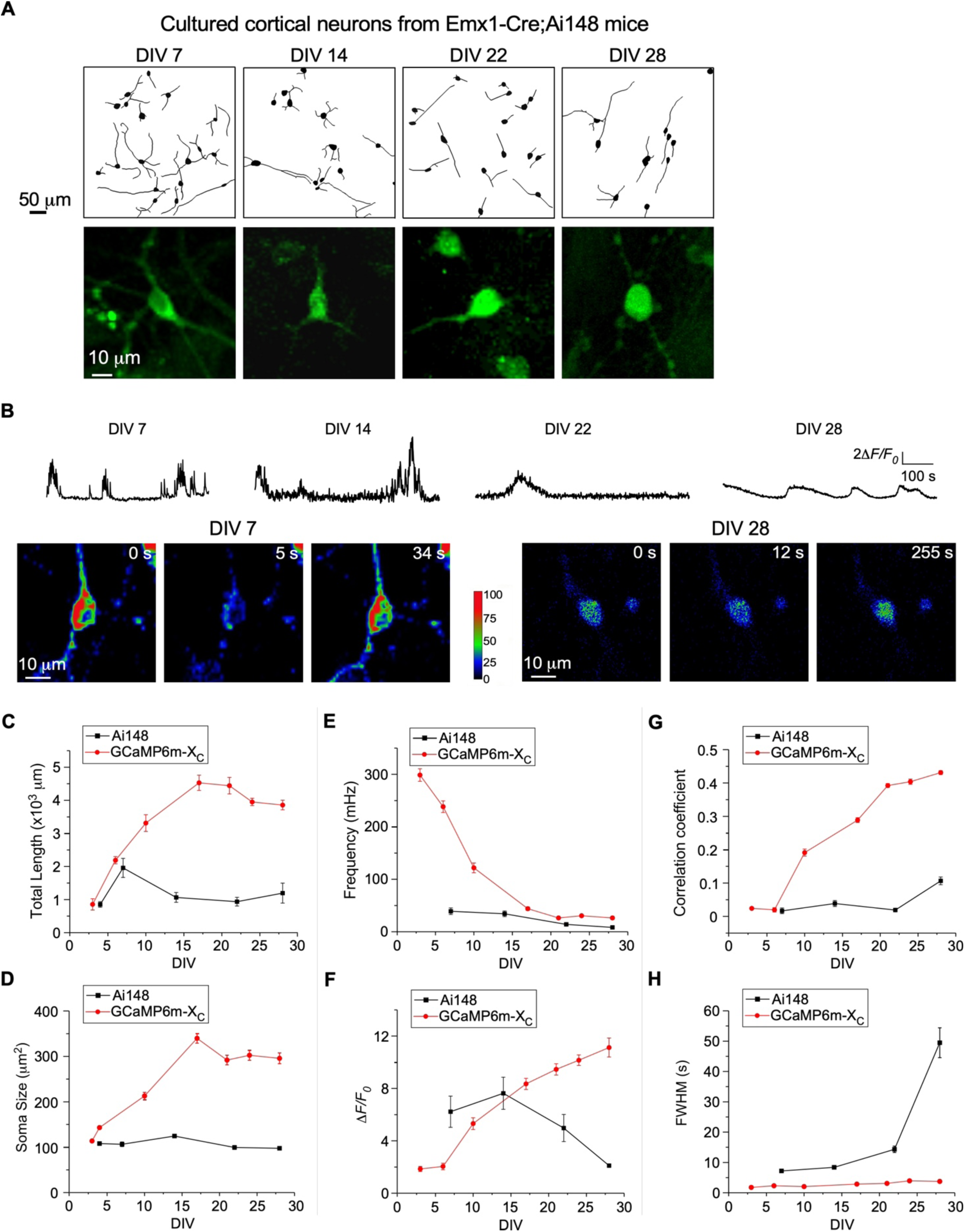
Long-term culturing and imaging of cortical neurons from Emx1-Cre;Ai148 mice with constitutive GCaMP6f expression. (**A**) Representative neurite tracing and fluorescence images for cultured cortical neurons constitutively expressing GCaMP6f from Emx1-Cre;Ai148 mice on selected DIV. (A) Oscillatory Ca^2+^ waveforms associated with color-coded fluorescence images. (**C** and **D**) Temporal profiles of total neurite length (**C**) and soma size (**D**). (**G-H**) Temporal profiles of oscillation frequency (mHz, **E**), peak amplitude (Δ*F/F0*, **F**), synchronization (correlation coefficient, **G**), and FWHM (sec, **H**). Red lines represent the average results (over 3 independent trials of ∼4-week culturing/imaging) from cortical neurons virally-expressing GCaMP6m-X_C_. Standard error of the mean (S.E.M) was calculated when applicable.

**Figure 3—video supplement 1.** Spontaneous Ca^2+^ oscillation of cultured cortical neurons virally-expressing GCaMP6m-X_C_ on DIV 6. Neurons at early stage of development exhibited high frequency, low amplitude and less synchronization. Video plays at 60x speed.

**Figure 3—video supplement 2.** Spontaneous Ca^2+^ oscillation of cultured cortical neurons virally-expressing GCaMP6m-X_C_ on DIV 10. Red areas indicate neurons with higher frequency and amplitude, whereas blue areas indicate neurons with relatively lower frequency and amplitude. Notably, neurons with strong Ca^2+^ oscillation were accompanied with well-developed neurites and connections. Video plays at 60x speed.

**Figure 3—video supplement 3.** Spontaneous Ca^2+^ oscillation of cultured cortical neurons virally-expressing GCaMP6m-X_C_ on DIV 28. Neurons exhibited highly synchronized Ca^2+^ oscillations with robust spikes and complicated neurite connections. Video plays at 60x speed.

**Figure 3—video supplement 4.** Spontaneous Ca^2+^ oscillation of cultured cortical neurons virally-expressing GCaMP6m on DIV 6. The Ca^2+^ activities of neurons were similar to neurons expressing GCaMP6m-X_C_, indicating that the damage of GCaMP was not significant at this stage. Video plays at 60x speed.

**Figure 3—video supplement 5.** Spontaneous Ca^2+^ oscillation of cultured cortical neurons virally-expressing GCaMP6m on DIV 10. Numbers of neurons exhibited broken neurites and soma. Few neurons with intact neurites still fired with robust Ca^2+^ spikes. Video plays at 60x speed.

**Figure 3—video supplement 6.** Spontaneous Ca^2+^ oscillation of cultured cortical neurons virally-expressing GCaMP6m on DIV 28. Most of the neurons were broken and no longer exhibited Ca^2+^ oscillation. Video plays at 60x speed.

**Figure 5—video supplement 1.** *In vivo* two-photon imaging of spontaneous Ca^2+^ oscillation of neurons virally-expressing GCaMP6m-X_C_ in S1 primary somatosensory cortex within optimal time window (4 weeks post virus). Images (512 x 512 pixels, 600 x 600 μm^2^) of L2/3 cells (150-250 μm under the pia) in S1BF were collected at 7.4 Hz frame rate. Video plays at 6x speed.

**Figure 5—video supplement 2.** *In vivo* two-photon imaging of spontaneous Ca^2+^ oscillation of neurons virally-expressing GCaMP6m-X_C_ in S1 primary somatosensory cortex without optimal time window (11 weeks post virus). Enlarged images (512 x 512 pixels, 300 x 300 μm^2^) of L2/3 cells (150-250 μm under the pia) in S1BF were collected at 7.4 Hz frame rate. Video plays at 6x speed.

**Figure 5—video supplement 3.** *In vivo* two-photon imaging of spontaneous Ca^2+^ oscillation of neurons virally-expressing GCaMP6m in S1 primary somatosensory cortex within optimal time window (4 weeks post virus). Enlarged images (512 x 512 pixels, 300 x 300 μm^2^) of L2/3 cells (150-250 μm under the pia) in S1BF were collected at 7.4 Hz frame rate. Neurons with nucleus-filled GCaMP6m barely fired. Video plays at 6x speed.

**Figure 5—video supplement 4.** *In vivo* two-photon imaging of spontaneous Ca^2+^ oscillation of neurons virally-expressing GCaMP6m in S1 primary somatosensory cortex without optimal time window (8 weeks post virus). Enlarged images (512 x 512 pixels, 300 x 300 μm^2^) of L2/3 cells (150-250 μm under the pia) in S1BF were collected at 7.4 Hz frame rate. Neurons with nucleus-filled GCaMP6m barely fired. Video plays at 6x speed.

**Figure 7—video supplement 1.** Spontaneous Ca^2+^ activities of cultured cortical neurons on DIV 7 from transgenic mice (Rasgrf2-2A-dCre x Ai148) for which GCaMP6f was induced by 10 μM TMP. Neurons at the early stage of development exhibited high frequency, low amplitude and less synchronization. Video plays at 60x speed.

**Figure 7—video supplement 2.** Autonomous Ca^2+^ oscillation of cultured cortical neurons on DIV 28 from transgenic mice (Rasgrf2-2A-dCre x Ai148) for which GCaMP6f was induced by 10 μM TMP. Broken neurites and damaged soma were evident in a large number of neurons. Few neurons still produced robust Ca^2+^ waveforms in coincidence with less damaged neurites. Video plays at 60x speed.

**Figure 8—video supplement 1.** Spontaneous Ca^2+^ oscillation of DIV-32 neurons co-expressing GCaMP6m-X_C_ and GCaMP6m-X_N_. Nuclear Ca^2+^ signals resemble cytosolic Ca^2+^ signals, but with much slower decay time. Video plays at 60x speed.

## Notes

### Competing Interest Statement

The authors have declared no competing interest.

